# Hair follicle-resident progenitor cells are a major cellular contributor to heterotopic subcutaneous ossifications in a mouse model of Albright hereditary osteodystrophy

**DOI:** 10.1101/2024.06.18.599506

**Authors:** Patrick McMullan, Peter Maye, Sierra H. Root, Qingfen Yang, Sarah Edie, David Rowe, Ivo Kalajzic, Emily L. Germain-Lee

**Affiliations:** Department of Pediatrics, University of Connecticut School of Medicine, Farmington, CT; Department of Reconstructive Sciences, Center for Regenerative Medicine and Skeletal Development, University of Connecticut School of Dental Medicine, Farmington, CT; The Jackson Laboratory, Farmington, CT; Albright Center, Division of Endocrinology & Diabetes, Connecticut Children’s, Farmington, CT

## Abstract

Heterotopic ossifications (HOs) are the pathologic process by which bone inappropriately forms outside of the skeletal system. Despite HOs being a persistent clinical problem in the general population, there are no definitive strategies for their prevention and treatment due to a limited understanding of the cellular and molecular mechanisms contributing to lesion development. One disease in which the development of heterotopic subcutaneous ossifications (SCOs) leads to morbidity is Albright hereditary osteodystrophy (AHO). AHO is caused by heterozygous inactivation of *GNAS*, the gene that encodes the α-stimulatory subunit (Gα_s_) of G proteins. Previously, we had shown using our laboratory’s AHO mouse model that SCOs develop around hair follicles (HFs). Here we show that SCO formation occurs due to inappropriate expansion and differentiation of HF-resident stem cells into osteoblasts. We also show in AHO patients and mice that *Secreted Frizzled Related Protein 2* (*SFRP2)* expression is upregulated in regions of SCO formation and that elimination of *Sfrp2* in male AHO mice exacerbates SCO development. These studies provide key insights into the cellular and molecular mechanisms contributing to SCO development and have implications for potential therapeutic modalities not only for AHO patients but also for patients suffering from HOs with other etiologies.

## Introduction

Heterotopic ossifications (HOs) are the result of a pathologic process by which bone inappropriately forms outside of the skeletal system in areas such as the dermis, subcutaneous tissue, and skeletal muscle [1–3] and are a significant clinical issue in the general population. HOs frequently form after surgical procedures such as hip arthroplasty (up to 40%), as well as close to 30% of fractures, high-energy military injuries, and severe burns, and up to 50% of traumatic brain and spinal cord injuries [1]. Although non-genetic forms of HOs are a major clinical issue, there are no definitive therapeutic strategies for their prevention and treatment. This lack of available therapies is problematic because HOs not only can cause pain and joint immobility but also can cause permanent neurologic and vascular insufficiency if inappropriately managed [1,2]. Furthermore, surgical resection of HOs is often not an option due to frequent post-operative recurrence [1,2].

A definitive strategy for the prevention and treatment of HOs requires a better understanding of the cellular populations contributing to their formation and the molecular mechanisms promoting aberrant osteogenesis. One approach towards understanding these etiologies is to study monogenic disorders that result in spontaneous heterotopic bone formation. To date, there are three monogenic disorders that are known to be characterized by the formation of extensive HOs. The most devastating in terms of HO severity is fibrodysplasia ossificans progressiva (FOP), which has been shown to be caused by mutations in the gene encoding activin A receptor type 1 (*ACVR1*) [4,5]. This mutation results in the hyperactive dysregulation of the BMP signaling cascade leading to endochondral bone formation within skeletal muscle and surrounding connective tissue [4,5]. Two additional disorders that result in spontaneous HO formation include Albright hereditary osteodystrophy (AHO) and progressive osseous heteroplasia (POH) (for review, [6–15]).

AHO is a disorder caused by the heterozygous inactivation of *GNAS*, an imprinted gene that encodes the α-stimulatory subunit (Gα_s_) of G protein-coupled receptors (GPCRs), which are utilized by multiple hormones that activate adenylyl cyclase [16,17]. Patients with maternally inherited *GNAS* mutations develop pseudohypoparathyroidism type 1A (PHP1A) and exhibit extraskeletal manifestations that include obesity and resistance to multiple hormones requiring Gα_s_, such as PTH, TSH, GHRH, and LH/FSH, [18–24] whereas patients with paternally derived *GNAS* mutations develop pseudopseudohypoparathyroidism (PPHP), in which patients have AHO skeletal features without severe obesity [25] or hormonal resistance (for review [6–11]). Through studies of both humans and mouse models [6] these metabolic and hormonal distinctions were shown to be due to tissue-specific paternal imprinting of *GNAS*, typically within endocrine organs such as the pituitary [22,23,26], thyroid [18–20,27], gonads [20,27], renal cortex [27–29] and potentially osteoclasts [30].

POH is also attributed to heterozygous inactivation of *GNAS*, secondary to paternal inheritance of the affected allele, and in general POH patients do not have hormonal resistance, [13–15,31] although there are rare cases of an overlap syndrome with PHP1A [32,33]. Unlike lesions in FOP, the HOs that form within AHO and POH develop within the skin and subcutaneous tissue by intramembranous ossification for which the cellular and molecular mechanisms remain undetermined [7,12,15,34,35]. Although AHO and POH share a similar genetic defect, they are recognized clinically as two distinct disorders aside from the difference in hormonal resistance (for review, [6–15]). First, the extent of penetration of heterotopic ossifications differs between the two conditions. In AHO, heterotopic bone formation is always restricted to the dermis and subcutaneous tissue and does not penetrate further [10,12,35]. Our group has confirmed this through physical examinations of our AHO patient population as well as through clinically-indicated radiographs, computerized tomography (CT) scans, and magnetic resonance images (MRI) [35], and this is recapitulated in our mouse model [34]. Therefore, heterotopic bone lesions that form in AHO are defined as subcutaneous ossifications (SCOs). Patients with POH, however, develop significantly more invasive heterotopic bone when compared to AHO, and although ossifications in POH can be identified within the dermis and subcutaneous tissue, these lesions often penetrate into underlying tissue such as skeletal muscle, fascia, tendons, and deep connective tissue [7–15,35]. The second distinction between AHO and POH is that in addition to heterotopic bone formation, AHO patients develop additional skeletal manifestations including adult short stature and brachydactyly, whereas these are typically absent in POH [6–15][22–24].

In our Albright Center, a clinic dedicated to the care of patients with AHO, we have evaluated hundreds of mutation-confirmed patients from throughout the world. Many suffer from pain and decreased functional abilities secondary to SCOs, and in this regard, we have become interested in determining the mechanisms involved in SCO initiation and formation; an understanding of this pathophysiology could provide insights into therapeutic modalities for prevention and treatment. Over a 16-year timespan of monitoring a cohort of mutation–confirmed AHO patients, we found that SCOs are present at an equal prevalence of approximately 70% in both PHP1A and PPHP [35]. These lesions either develop *de novo* as early as birth or secondary to repetitive pressure or trauma. Further observation of this patient cohort revealed that SCO prevalence was significantly higher among male patients than females, suggesting the potential for sex hormones contributing to SCO development. Additionally, patients with nonsense or frameshift *GNAS* mutations developed SCOs at a significantly higher frequency (>90%) than patients with missense mutations (29.2%), suggesting a genotype-phenotype correlation [35].

In conjunction with clinically monitoring AHO patients, we had generated and characterized an AHO mouse model via the targeted disruption of exon 1 of *Gnas* that recapitulates the human disorder [27,34]. In particular, mice with heterozygous inactivation of *Gnas* (*Gnas E1+/-*) [27] develop SCOs that are independent of the parental origin of the mutant allele [34]*. Gnas E1+/-* mice form SCOs spontaneously and/or in response to repetitive pressure or trauma, such as at the base of the tail, footpads, and surrounding ear tags, and CT examination of *Gnas E1+/-* mice revealed that SCOs are limited to the dermis and subcutaneous tissue [34]. Similar to AHO patients, male *Gnas E1+/-* mice have SCOs at a significantly higher prevalence than female mice, with 100% of male mice developing SCOs by 9 months of age. Histologic evaluation revealed that prior to the formation of radiographically-detectable SCOs, male *Gnas E1+/-* mice at 3 months exhibit hypercellularity and collagen deposition within the reticular dermis that specifically surrounds hair follicles (HF) [34]. These histologic changes appear to be essential for the future development of SCOs since both male and female *Gnas E1+/-* mice at later timepoints develop SCOs that consistently form directly adjacent to or surrounding HFs. The HF is known to contain both epithelial and mesenchymal-derived progenitor populations that maintain their proliferative abilities throughout all phases of life [36–38], and this consistent spatial localization of SCOs near HFs suggested the possibility that this microenvironment and its progenitor populations may play a role in ossification development. This investigation is the first to demonstrate that SCO formation is initiated by the inappropriate expansion and differentiation of HF-resident dermal sheath cells into osteoblasts.

## Results

### Secreted Frizzled Related Protein 2 (*SFRP2)* expression correlates with SCO severity in human fibroblasts isolated from AHO and POH participant skin biopsies

As an initial investigation to identify genes and/or pathways that may play a role in SCO formation, we had performed a microarray analysis on RNA from primary dermal fibroblast cultures generated from skin biopsies from a selected group of mutation-confirmed AHO and POH participants. The participants selected either lacked SCOs or had SCOs of varying severity: Participant 1 (P1) was an adult female with PPHP with moderate palpable SCOs; Participant 2 (P2) was an adult female with PPHP without palpable SCOs; Participant 3 (P3) was a female child, daughter of P2, with PHP1A and severe SCOs; Participant 4 (P4) was an adult male with POH and extensive ossifications in the subcutaneous and deep connective tissue that invaded into the muscle, nerve, and blood vessels as documented by both surgical pathology of excised ossifications and imaging performed for clinical reasons (CT and/or MRI). Participant 5 (P5) was an adolescent male with PPHP and one small palpable SCO. We selected this diverse subgroup for analysis to examine both pre- and post-pubertal males and females with PHP1A and PPHP, and differences in the extent of SCO formation between close relatives.

We assessed differential gene expression by performing three comparisons (Figure 1A) that we hypothesized would have the greatest potential of highlighting variations in expression based on the degree of SCO formation and that would also help sort out potential hormonal effects that could be leading to SCOs being worse in males than in females based on our past studies in both humans and our mouse model [34,35]. In all three comparisons (P1 vs P2, P3 vs P2, and P4 vs P5), a participant with the greater number (as well as size) of SCOs was compared to a participant with fewer or no ossifications. We identified 23 differentially regulated genes (14 upregulated and 9 downregulated) that were observed within SCO-containing regions in each comparison (Figure 1B,C). The two most upregulated genes were Rho GTPase Activating Protein 28 (*ARHGAP28*), and Secreted Frizzled Related Protein 2 (*SFRP2*), which have been shown to become activated in response to extracellular matrix assembly and negatively regulate stress fiber formation both *in vivo* and *in vitro* [39–43]. Microarray analysis also revealed an upregulation in *ALDH1A3* and *NR4A3*, which are genes that have since been identified as being commonly recognized biomarkers for carcinoma-associated fibroblasts (CAFs) in skin disorders including basal cell carcinoma and systemic sclerosis [44–47].

**Figure 1:**
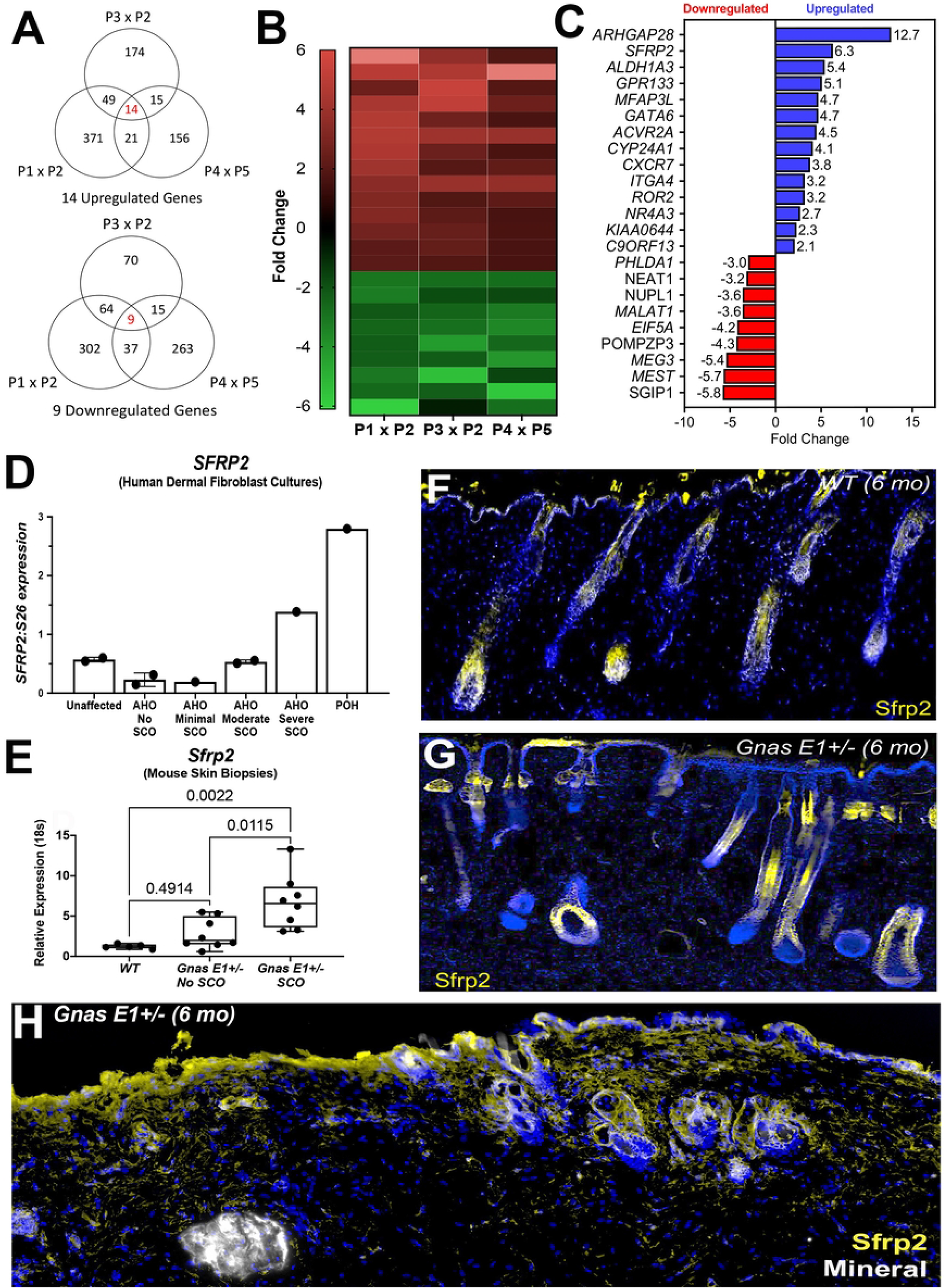
Microarray analyses of AHO/POH human dermal fibroblasts identify transcriptional profiles of fibroblasts in SCO-containing regions and are consistent with murine profiles. (A) Venn diagram of differentially regulated genes between three microarray analyses from RNA isolated from AHO and POH human dermal fibroblast cultures. (B) Representative heatmap (C) Listing of commonly differentially regulated genes among the three microarray comparisons. (D) Quantification of Northern blot data of *SFRP2* expression relative to S26 from AHO and POH human dermal fibroblasts. (E) RT-PCR of *Sfrp2* expression for 12-month WT and *Gnas* E1+/-skin samples. (F-G) Dorsal skin of 6-month-old (F) WT and (G) unaffected *Gnas* E1+/- mice stained with anti-Sfrp2 (yellow). (H) Dorsal skin section of 6-month *Gnas* E1+/- mouse SCO skin stained with anti-Sfrp2 (yellow).

Among these differentially expressed genes, we became most interested in further examining the role of *SFRP2* in SCO pathogenesis given that it was the most upregulated gene with a known relationship to osteogenesis at the time of microarray investigation [39,48]. Additionally, we had identified a direct correlation between *SFRP2* mRNA expression and SCO severity by Northern blot analysis of RNA isolated from skin biopsies from 7 participants in our investigations who had either AHO or POH or 2 who were unaffected family members (described in methods) (Figure 1D). In particular, we found the highest level of *SFRP2* mRNA expression in the POH participant, intermediate expression in the AHO participant with severe ossifications, and lower levels of expression in the remaining participants with minimal, moderate, or no ossifications.

### *Gnas E1+/-* mice display elevated *Sfrp2* expression within the hair follicle microenvironment surrounding ossification sites

To further explore the functional role of *SFRP2* in the development of SCOs, we utilized our *Gnas E1+/-* mouse model that phenotypically recapitulates lesion development [27,34]. We assessed whether the transcriptional differences observed in human samples were similar to those within skin samples harvested from 12-month-old *Gnas E1+/-* mice (Figure 1E, Supplemental Figure 1). *Gnas E1+/-* mice displayed a significant upregulation of *Sfrp2* mRNA expression in dorsal skin samples containing SCOs when compared to both *WT* skin samples and *Gnas E1+/-* samples harvested from unaffected skin regions (Figure 1E). We also confirmed that the upregulation of *ARHGAP28, ALDH1A3, GPR133, ACVR2A* and *ROR2* observed in human SCO samples was similarly observed in *Gnas E1+/-* SCO (Supplemental Figure 1), demonstrating further correlation of our mouse model with the human disorder.

Based on this direct correlation of *SFRP2* expression and SCO severity in both the human and mouse samples, we assessed the spatial localization of Sfrp2 protein within the dorsal skin of 6-month *WT* and *Gnas E1+/-* mice by immunofluorescence (Figure 1F-H). Sfrp2 expression was observed in epithelial-derived cells in the HF and in a limited number of dermal fibroblasts (Figure 1F,G). However, *Gnas E1+/-* SCO-containing skin samples revealed a markedly different pattern (Figure 1H) and showed an expansion of Sfrp2+ cellular populations along the basal epithelial surface and surrounding HFs. We detected limited expression of *Sfrp2* in cells on the SCO bone-lining surface (Figure 1H) and observed a similar pattern of Sfrp2+ cells in dorsal skin samples from 15-month *Gnas E1+/-* mice with extensive SCOs (Supplemental Figure 2).

Given that SCO-containing skin regions exhibit a broader expression of *Sfrp2* near HFs and that our previous studies had found SCOs consistently developing adjacent to or surrounding HFs, we further examined the contribution of HF cellular populations to SCO formation and how *Sfrp2* may influence the differentiation capacity of cell types within this microenvironment.

### *Gnas E1+/-* mice form SCOs that progressively expand and localize to hair follicles

We next analyzed skin samples of *WT* and *Gnas E1+/-* mice by radiographic imaging and histology (Figure 2). Histologic analysis of *Gnas E1+/-* mice demonstrated that SCO formation occurs through intramembranous ossification as indicated by enhanced collagen and osteoid deposition along the bone lining surface by Masson Trichrome staining and the absence of glycosaminoglycan detection by Safranin O staining (Supplemental Figure 3). Serial x-rays of *Gnas E1+/-* mice demonstrated that SCOs become radiographically detectable by 4 months of age and progressively expand (Figure 2A). We have found that this progressive expansion of intramembranous heterotopic bone surrounding HFs in *Gnas E1+/-* mice is reflected by the presence of both actively mineralizing osteoblasts (Figure 2B) and bone-lining osteoclasts (Supplemental Figure 3). We therefore assessed the spatial localization of progenitor cells within the HF by alkaline phosphatase (ALP) histochemistry in dorsal skin sections of 15-month-old mice (Figure 2C-F). Both *WT* (Figure 2C) and *Gnas E1+/-* (Figure 2D) mice displayed ALP+ populations within the HF that localized to a distinct mesenchymal population, specifically the dermal papilla. However, *Gnas E1+/-* mice also exhibited an expansion of ALP+ cells within two additional areas of the dermis in SCO regions (Figure 2D-F), which included the SCO bone-lining surface and the adjacent unmineralized regions of the dermis encompassing the entire HF.

**Figure 2:**
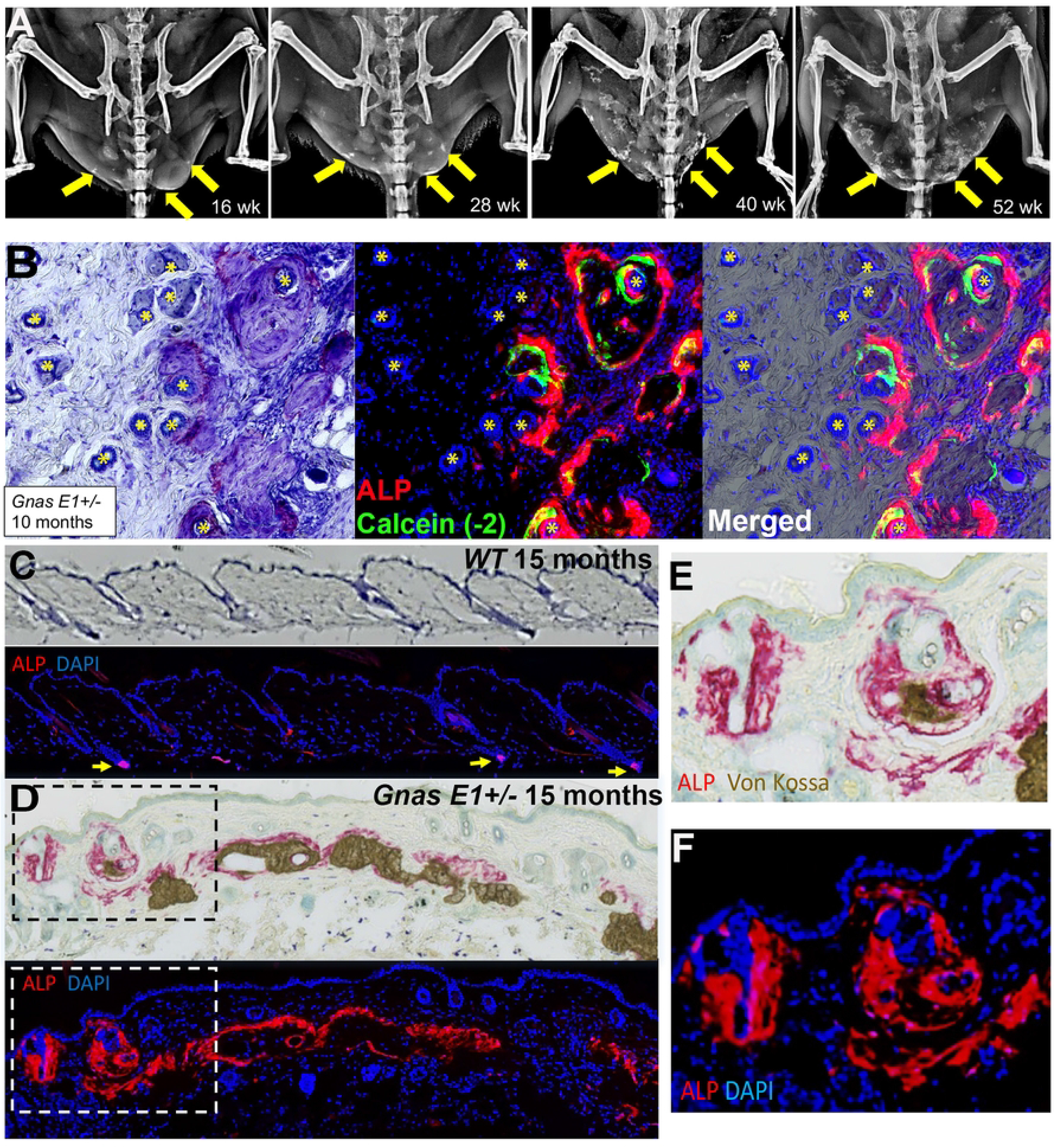
*Gnas E1+/-* mice develop progressively expanding subcutaneous ossifications in dermis surrounding hair follicles. (A) Consecutive x-ray images of a male *Gnas E1+/-* mouse at 16, 28, 40, and 52 weeks. (B) Dorsal skin of 40-week-old male *Gnas E1+/-* mouse demonstrating SCOs surrounding hair follicles and the presence of active mineralizing osteoblasts [alkaline phosphatase (ALP+) populations superimposed over a calcein mineralization label]. (C-D) Dorsal skin sections from 15-month-old (C) *WT* and (D) *Gnas E1+/-* mice stained using Toluidine Blue, ALP, and Von Kossa. ALP+ populations within *WT* mice are limited to the dermal papilla (yellow arrows) whereas *Gnas E1+/-* mice display ALP+ cells within the dermal papilla and throughout the dermis along the SCO bone surface.(E-F). Higher power images of boxed regions in panel D.

### *Osterix-mCherry* reporter identifies expanded hair follicle progenitor cells that are osteoprecursors within *Gnas E1+/-* mice as αSMA+ dermal sheath cells *in vivo*

We next crossed *Gnas E1+/-* mice with *Osterix-mCherry* (*Osx-mCherry*) transgenic reporter mice (Figure 3A) in order to label multipotent mesenchymal progenitors, osteoblasts, and osteocytes *in vivo* [49]. We were particularly interested in examining skin regions prior to the formation of radiographically detectable SCOs based on our previous studies demonstrating hypercellularity and collagen deposition near HFs [34]. We hypothesized that this hypercellularity represented the condensation of differentiated HF osteoprecursors initiating intramembranous ossification.

**Figure 3:**
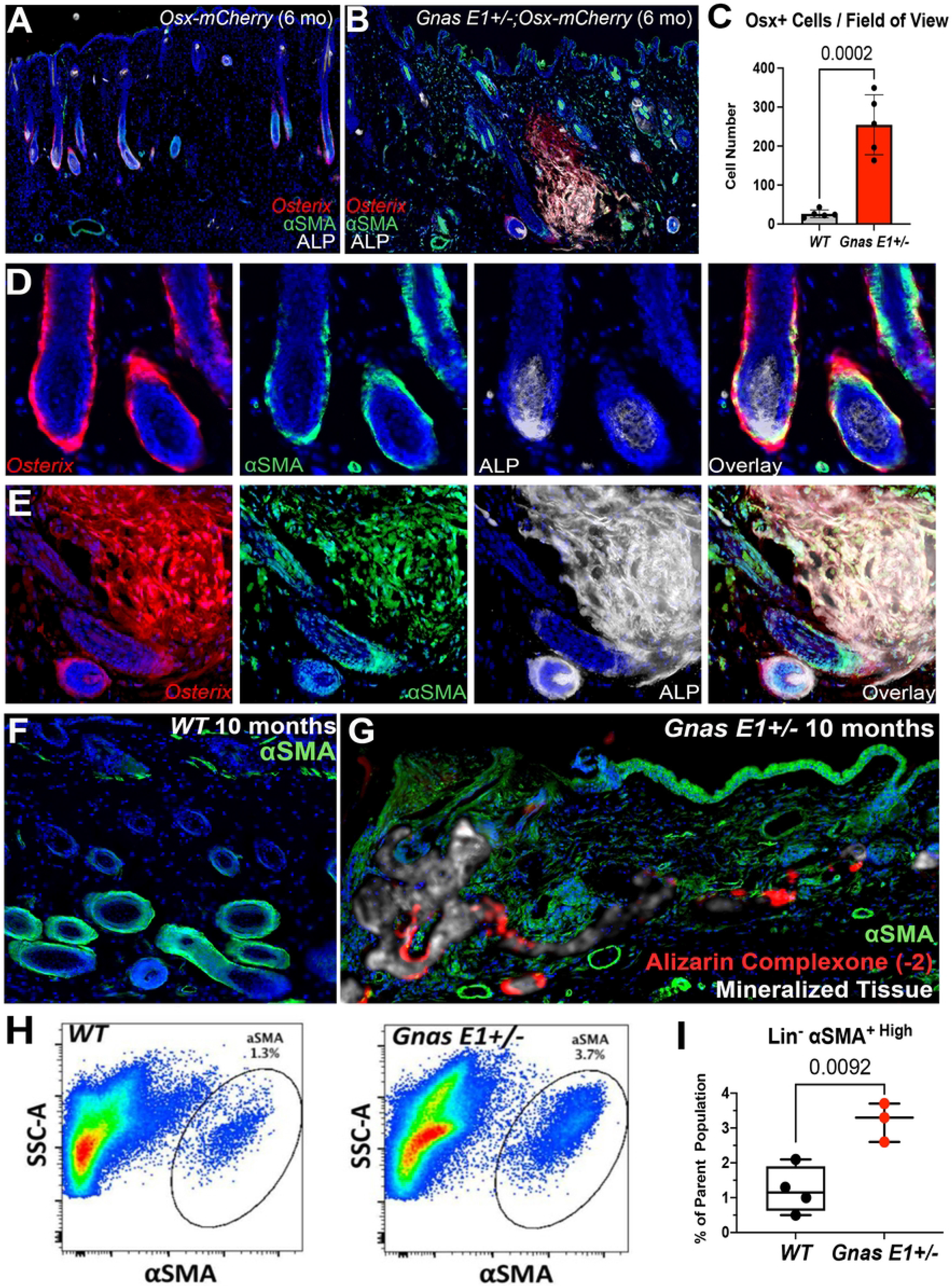
Genetic fate-mapping studies using *Osterix-mCherry* model identifies hair-follicle resident osteoprecursors as αSMA+ dermal sheath cells (A-B) Dorsal skin sections of 6-month. (A) *Osx-mCherry* and (B) *Gnas E1+/-;Osx-mCherry* littermates demonstrating spatial expression of *Osterix* (red), alpha-Smooth Muscle Actin (αSMA) (green), and ALP (white). (C) Bar graph demonstrating number of *Osterix+* cell types within the dermis of 6-month *Osx-mCherry* and *Gnas E1+/-;Osx-mCherry* mice. (D-E) Higher-power images of *Osx-mCherry* and *Gnas E1+/-;Osx-mCherry* hair follicles demonstrating *Osterix* and αSMA colocalization. (F-G) Representative dorsal skin sections of 10-month-old (F) *WT* and (G) *Gnas E1+/-* mice stained for αSMA (green), mineralized tissue (white), and bone mineral label alizarin complexone (red). (H) Representative flow cytometry plots. SSC-A refers to side scatter area. (I) Graph of the percentage of Lin-αSMA+ populations from 10-month-old *WT* and *Gnas E1+/-* mice *in vivo*.

Histologic evaluation at 3 weeks (Supplemental Figure 4) and 6 months of age (Figure 3A-E) identified a distinct population of Osterix+ cells in *Gnas E1+/-* mice that localized to the outer surface of the HF. These Osterix+ cells aligned with the location of the expanded ALP+ osteprecursors that we had previously identified within *Gnas E1+/-* skin samples (Figure 2D-F). These Osterix+ populations corresponded to dermal sheath cells based upon their co-expression of alpha-Smooth Muscle Actin (αSMA), which is an established biomarker for dermal sheath cells (Figure 3D) [50,51]. It is important to note that although αSMA has been shown to label dermal sheath cells, it has also been shown to label smooth muscle cells within the HF arrector pili muscle and the underlying blood vasculature (Figure 3A,B) [50,51]. We did not detect Osterix expression in dermal papilla cells (Figure 3C). Given that the only Osterix+ αSMA+ double positive populations within the dermis localized to the dermal sheath, we hypothesized that dermal sheath cells may contribute to the process of SCO initiation.

When evaluating 6-month *Gnas E1+/-;Osx-mCherry* mice in skin regions without radiographically detectable SCOs (Figure 3C, 3E), we observed a significant expansion of Osterix+ cell populations that extended into the dermis from adjacent HFs when compared to littermate *Osx-mCherry* mice. Immunofluorescence colocalization studies revealed that these expanded Osterix+ cells co-expressed αSMA and ALP (Figure 3E). These data aligned with our initial hypothesis and suggested that these Osterix+, αSMA+, ALP+ triple positive cells within the dermis are labeling expanded HF-derived osteoprecursors that are actively undergoing osteogenic differentiation. Additionally, these data demonstrating Osterix and αSMA colocalization were of particular interest because αSMA has been previously identified as a biomarker for tissue-resident mesenchymal progenitors with osteogenic potential and have been shown to contribute to the initiation and progression of HOs within skeletal muscle [52–55]. We next evaluated the spatial localization of αSMA+ populations in 10-month-old mice (Figure 3F,G) because *Gnas E1+/-* mice at this timepoint consistently exhibit extensive SCOs throughout the dorsal skin. αSMA immunofluorescence in both *WT* and unaffected *Gnas E1+/-* skin regions labeled the dermal sheath cells as well as the HF arrector pili muscle and underlying vasculature, as similarly observed at earlier timepoints (Figure 3F). However, *Gnas E1+/-* mice exhibited an expansion of αSMA+ populations throughout the dermis that localized to areas of active bone mineralization, as indicated by alizarin complexone labeling (Figure 3G). In addition to histologic analysis, we also performed flow cytometry on single cell suspensions of enzymatically digested dorsal skin samples from 10-month *WT* and *Gnas E1+/-* mice to determine the percentage of αSMA+ mesenchymal progenitors as defined by percentage of αSMA+ cells within the Lineage-gate, which excluded hematopoietic and endothelial cells by CD45-Ter119- and CD31-staining, respectively. We found that *Gnas E1+/-* mice exhibited a 3-fold increase in the percentage of αSMA+ mesenchymal progenitors when compared to *WT* samples (Figure 3H-I, Supplemental Figure 5). In summary, these data suggest that HF-resident αSMA+ mesenchymal populations may contribute to SCO development.

### Lineage tracing identifies osteoblasts and osteocytes in subcutaneous ossifications in *Gnas E1+/- ;αSMA^CreERT2^;Ai9^fl/fl^* mice

We next performed genetic fate-mapping studies by crossing our *Gnas E1+/-* mice with tamoxifen-inducible *⍺SMACre^ERT2^;Ai9^fl/fl^* mice (Figure 4A,B) [53,54] which would allow us to trace αSMA+ cell types at various timepoints in the skin and subcutaneous tissue following the injection of tamoxifen through the expression of Ai9 (tdTomato). We first examined the dorsal skin of 10-month-old *Gnas E1+/- ;⍺SMACre^ERT2^;Ai9^fl/fl^* and *⍺SMACre^ERT2^;Ai9^fl/fl^* mice that did not receive any tamoxifen in order to identify the degree of endogenous Ai9 expression due to leakiness, which was minimal in unaffected and SCO skin regions (Supplemental Figure 6).

**Figure 4:**
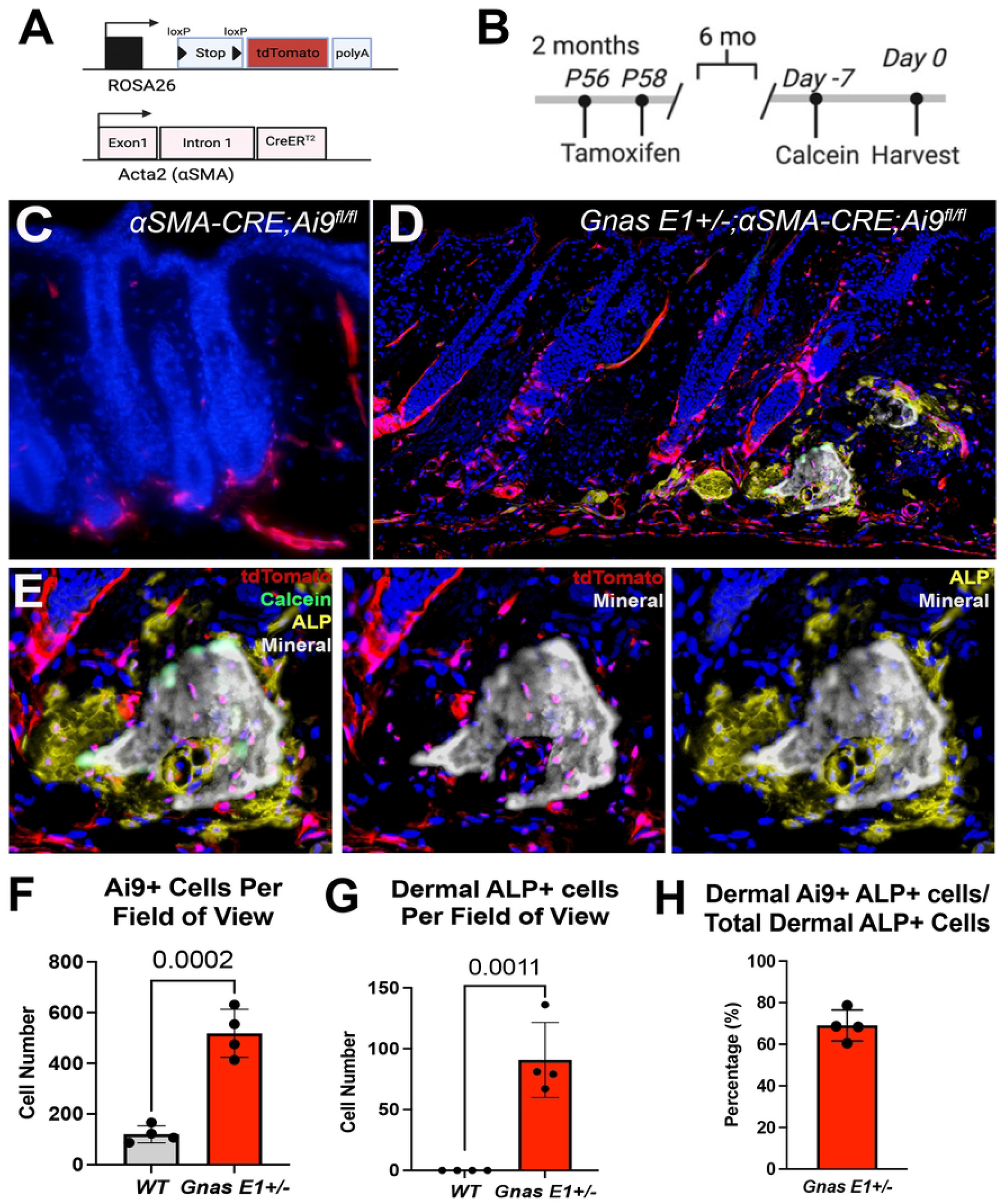
*In vivo* lineage tracing of αSMA+ populations identifies progenitor cells that are osteoprecursors contributing to SCO initiation. (A) Breeding scheme and generation of *Gnas E1+/- ;⍺SMACre^ERT2^;Ai9^fl/fl^* mice. (B) *In vivo* lineage tracing strategies utilized. (C-D) Dorsal skin sections of 8-month old mice: (C) *⍺SMACre^ERT2^;Ai9^fl/fl^* and (D) *Gnas E1+/-;⍺SMACre^ERT2^;Ai9^fl/fl^* following 6-month lineage tracing study of Ai9+ populations also shown at higher magnification in (E) in which *Gnas E1+/- ;⍺SMACre^ERT2^;Ai9^fl/fl^* mice displayed an expansion of Ai9+ populations near hair follicles and differentiated into SCO-lining osteoblasts (ALP+ cells over calcein label) and osteocytes embedded into SCO bone matrix. (F-G) Graph of number of dermal (F) Ai9+ and (G) dermal ALP+ populations within 8-month old *⍺SMACre^ERT2^;Ai9^fl/fl^* and *Gnas E1+/-;⍺SMACre^ERT2^;Ai9^fl/fl^* mice. (H) Graph demonstrating the ratio of dermal Ai9+;ALP+ double positive cells when compared to the total number of dermal ALP+ cells within 8-month *Gnas E1+/-;⍺SMACre^ERT2^;Ai9^fl/fl^* mice.

In alignment with previous αSMA lineage tracing studies in the skin [50,56]. we observed that 2 days following tamoxifen administration, Ai9 expression localized to the dermal sheath, HF arrector pili muscle, and underlying blood vasculature (Supplemental Figure 7). We next addressed the question of whether αSMA+ populations were essential cell types for the initiation of SCO formation (Figure 4C-H) by examining mice 6 months following tamoxifen treatment (Figure 4A-B for schematic). Histology of the dorsal skin from 8-month-old *⍺SMACre^ERT2^;Ai9^fl/fl^* mice showed that Ai9+ populations remained restricted to the dermal sheath, arrector pili, and blood vasculature (Figure 4C). However, within the surrounding HF microenvironment, *Gnas E1+/-;⍺SMACre^ERT2^;Ai9^fl/fl^* mice displayed a significant expansion of both Ai9+ and ALP+ populations (Figure 4D-G). Immunofluorescence colocalization studies revealed that 69% of these expanded Ai9+ populations within *Gnas E1+/-;⍺SMACre^ERT2^;Ai9^fl/fl^* mice coexpressed ALP (Figure 4H). Further colocalization analysis revealed these ALP+ Ai9+ double positive populations were localized to the SCO bone-lining surface superimposed over a calcein mineral label, and were embedded within the SCO bone matrix. Taken together, these data demonstrate that HF-resident αSMA+ cells are capable of undergoing differentiation into both osteoblasts and osteocytes and appear to be an essential cell type required for the initial osteoid deposition and formation of SCOs.

We next assessed the contribution of αSMA+ populations in the progressive expansion of SCOs by performing 21-day tamoxifen pulse-chase experiments in 9-month-old *Gnas E1+/-;⍺SMACre^ERT2^;Ai9^fl/fl^* mice with radiographically detectable SCOs compared to littermate *⍺SMACre^ERT2^;Ai9^fl/fl^* mice (Figure 5A). We did not observe any significant changes in the localization of Ai9+ populations in 9-month-old *⍺SMACre^ERT2^;Ai9^fl/fl^* mice (Figure 5B,C) when compared to earlier timepoints. We did, however, observe an expansion of Ai9+ populations surrounding HFs in the dermis of *Gnas E1+/-;⍺SMACre^ERT2^;Ai9^fl/fl^* mice within both unaffected and SCO skin regions (Figure 5D-G). Furthermore, *Gnas E1+/-;⍺SMACre^ERT2^;Ai9^fl/fl^* mice displayed Ai9+ populations along the SCO bone lining surface that: (1) expressed ALP superimposed over a calcein mineral label (Figure 5H); (2) expressed Sclerostin (Sost); and (3) were embedded within the SCO bone matrix (Figure 5H). These data further underscore that αSMA+ dermal sheath cells in *Gnas E1+/-* mice contribute not only to the initiation of SCO formation through expansion within the HF microenvironment but also serve as essential cells in SCO expansion given their osteogenic capacity and potential to differentiate into osteoblasts and osteocytes.

**Figure 5:**
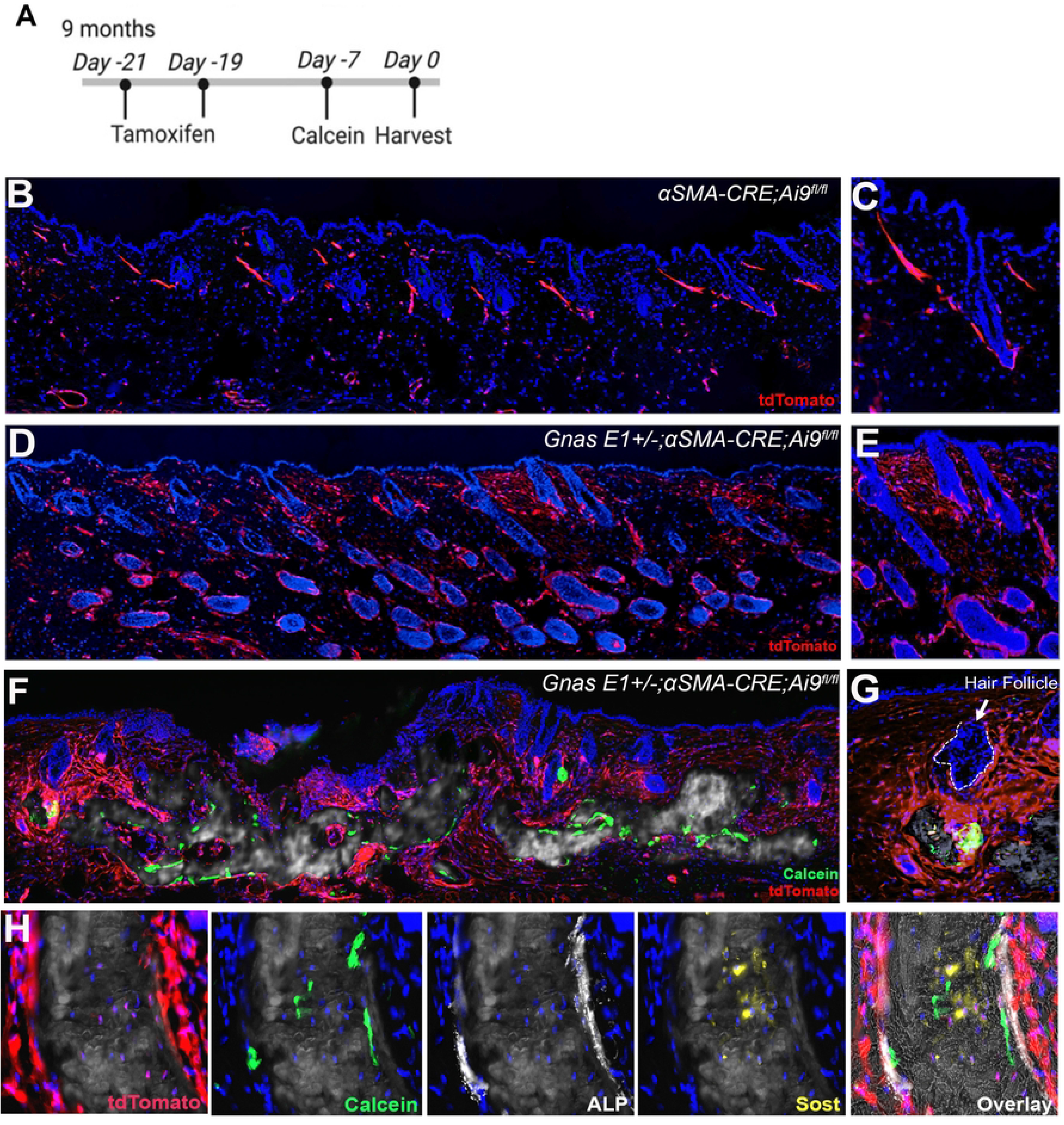
*In vivo* lineage tracing of αSMA+ populations identifies osteoprecursors contributing to SCO progression. (A) *In vivo* lineage tracing strategy utilized. (B-G) Dorsal skin sections of 10-month (B-C) *⍺SMACre^ERT2^;Ai9^fl/fl^* and (D-G) *Gnas E1+/-;⍺SMACre^ERT2^;Ai9^fl/fl^* mice within (D-E) unaffected and (F-G) SCO skin regions following 21-day lineage traceing. (H) SCO region of 10-month old *Gnas E1+/- ;⍺SMACre^ERT2^;Ai9^fl/fl^* mice demonstrating the differentiation of Ai9+ cells into bone lining osteoblasts and osteocytes (based on Sclerostin [Sost] coexpression).

In conjunction with *in vivo* lineage tracing, we assessed the osteogenic capacity of labeled Ai9+ populations *in vitro* by injecting 9-month-old *Gnas E1+/-;⍺SMACre^ERT2^;Ai9^fl/fl^* and *⍺SMACre^ERT2^;Ai9^fl/fl^* mice with tamoxifen at 7 and at 5 days prior to tissue harvest and then generating primary dermal explant cell cultures (Methods described[57]). This culture model allowed us to monitor the behavior of Ai9+ cells from the onset of culture establishment and throughout the course of our *in vitro* analyses (Figure 6A,B) using live cell imaging. Following 14 days of primary culture expansion, we performed FACS sorting analyses and confirmed a 3-fold increase in the percentage of Ai9+ populations in *Gnas E1+/-;⍺SMACre^ERT2^;Ai9^fl/fl^* cultures compared to *WT ⍺SMACre^ERT2^;Ai9^fl/fl^* (Figure 6C-D, Supplemental Figure 8). Gene expression studies demonstrated that Ai9+ sorted populations from *Gnas E1+/-* cultures have significant upregulation of *Osterix* (*Sp7*) mRNA expression when compared to *WT* Ai9+, *Gnas E1+/-* and *WT* unsorted cultures (Figure 6E). We next treated dermal explant cultures for 4 weeks with osteogenic induction media (Figure 6F-H). *Gnas E1+/-* cultures displayed multiple colonies of ALP+ populations and mineral deposition by Von Kossa staining (Figure 6G). Live imaging studies of *Gnas E1+/-;⍺SMACre^ERT2^;Ai9^fl/fl^* cultures revealed that *in vivo* labeled Ai9+ populations localized to areas of active mineralization based upon calcein staining (Figure 6G). This enhanced mineralization capacity in *Gnas E1+/-;⍺SMACre^ERT2^;Ai9^fl/fl^* cultures correlated with an upregulation in both *Osterix (Sp7)* and *Integrin Binding Sialoprotein* (*Ibsp*) mRNA expression by RT-PCR when compared to *⍺SMACre^ERT2^;Ai9^fl/fl^* cultures (Figure 6H). These data further implicate dermal αSMA+ progenitors as an essential cell type in SCO formation.

**Figure 6:**
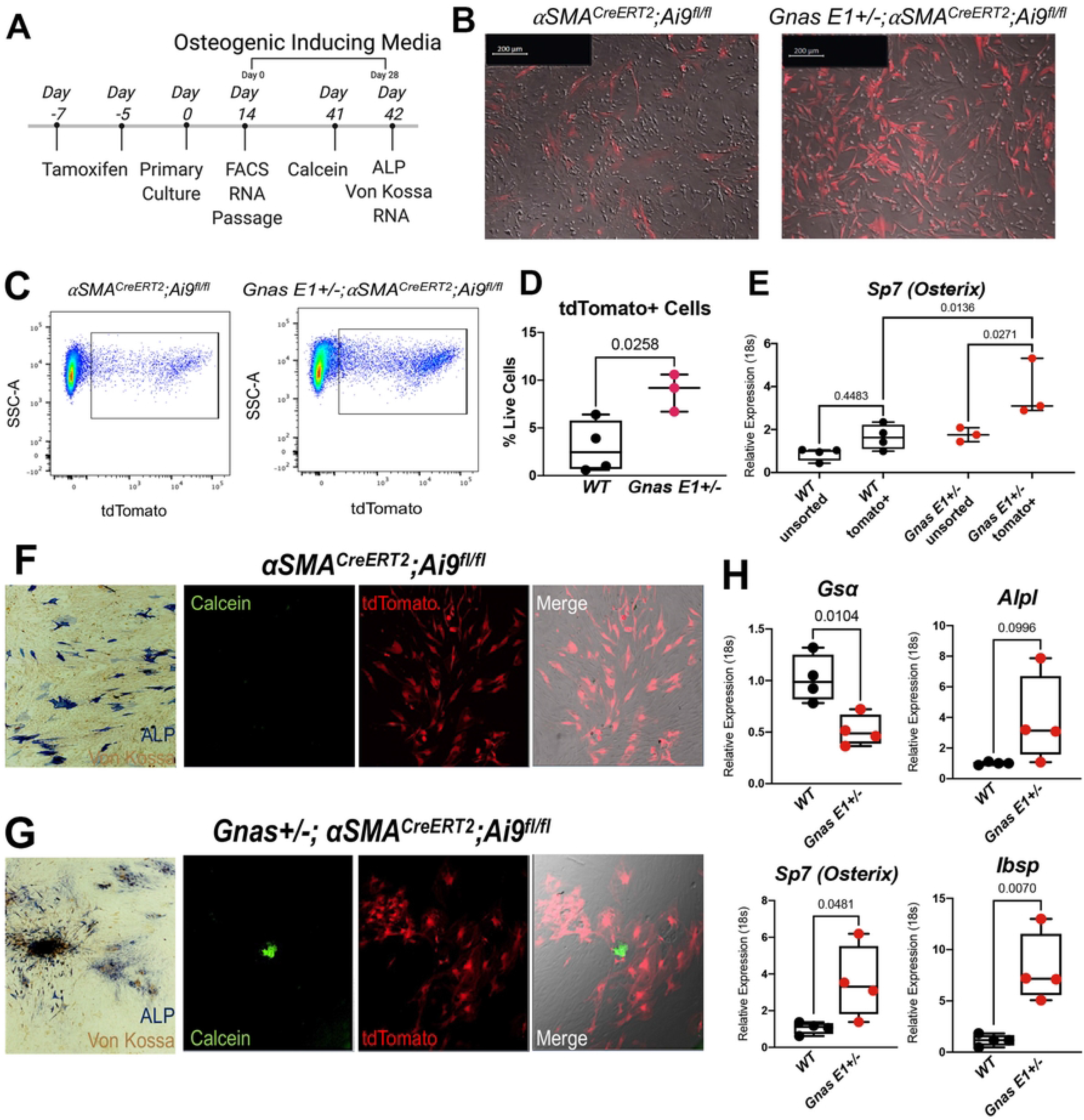
*In vivo* labeled Ai9+ populations from *Gnas E1+/-;⍺SMACre^ERT2^;Ai9^fl/fl^* mice exhibit enhanced osteogenic capacity *in vitro*. (A) Timeline of dermal explant culture generation and *in vitro* lineage tracing. (B) Representative images of *in vivo* labeled Ai9+ cells within primary dermal explant cultures. (C) FACS plots and (D) graph of the percentage of Ai9+ cells isolated from *⍺SMACre^ERT2^;Ai9^fl/fl^* and *Gnas E1+/-;⍺SMACre^ERT2^;Ai9^fl/fl^* cultures. (E) RT-PCR of *Sp7* (*Osterix*) mRNA expression for unsorted and Ai9+ sorted primary dermal explants. (F) *⍺SMACre^ERT2^;Ai9^fl/fl^* and (G) *Gnas E1+/-;⍺SMACre^ERT2^;Ai9^fl/fl^* dermal explant cultures following 28 days of osteogenic differentiation stained for ALP andVon Kossa as well as live culture images of calcein (green) and tdTomato (red). (H) RT-PCR of *Gs⍺, Alpl, Sp7* and *Ibsp* mRNA expression in cultures following 28 days of osteogenic differentiation.

### *Gnas E1+/-* mice exhibit transcriptome variation within SCO containing regions

We next investigated potential signaling pathways that may be promoting the inappropriate differentiation of these dermal populations by isolating RNA from dorsal skin samples from 12-month *WT* mice and *Gnas E1+/-* mice from both unaffected and SCO-containing skin regions and assessed transcriptional variations among samples using an RT-PCR array (Figure 7A, Supplemental Figure 9). We were particularly focused on a panel of genes related to Wnt, Sonic hedgehog (Shh), TGF-β, and BMP signaling pathways due to their implications in regulating both epithelial-mesenchymal interactions within the HF microenvironment [58–62] and the pathogenesis of heterotopic ossifications. We did not identify any significant differences in mRNA expression patterns between *WT* and unaffected *Gnas E1+/-* skin samples, but we observed significant transcriptional changes in genes related to each of these canonical signaling pathways within SCO-containing *Gnas E1+/-* skin samples when compared to both *WT* and unaffected *Gnas E1+/-* samples (Figure 7A, Supplemental Figure 9). We also confirmed the validity of these transcriptomic data by performing immunofluorescence within dorsal skin specimens from 12 month *WT* and *Gnas E1+/-* mice (Figure 7B-E), which demonstrated increased expression of Gli1+ cellular populations (Figure 7B,C) in addition to Tgfβ1+ populations (Figure 7D,E) throughout the basal epithelium, dermis, and SCO bone-lining surface of *Gnas E1+/-* skin samples when compared to *WT.* In summary, these findings suggest that *Gnas* heterozygous inactivation results in the dysregulation of multiple signaling pathways within the HF microenvironment to promote the osteogenic differentiating capacity of tissue-resident progenitor populations during SCO formation.

**Figure 7:**
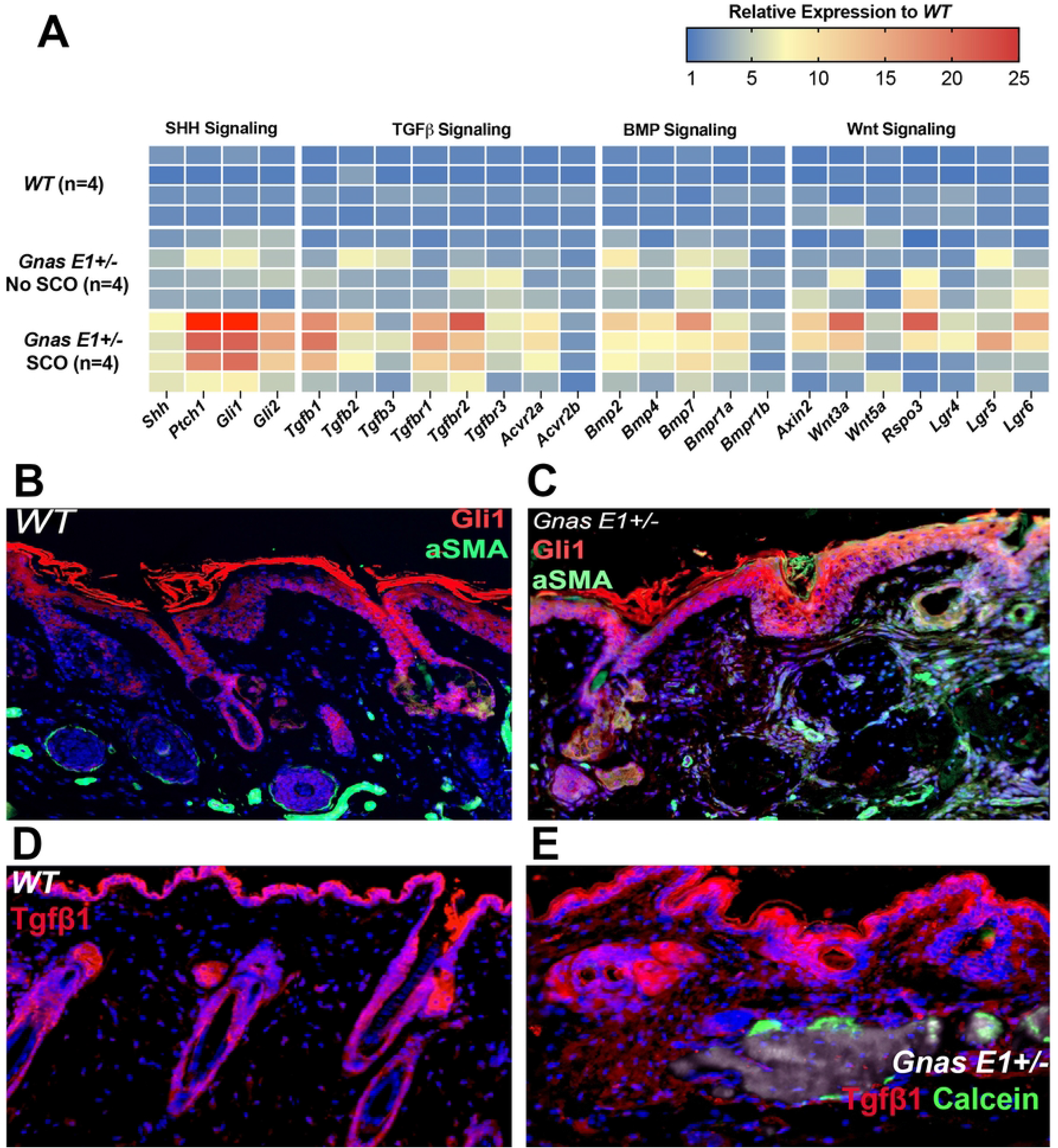
*Gnas E1+/-* mice exhibit transcriptome variation within SCO-containing regions. (A) Heatmap of differential gene expression in *Gnas* E1+/- SCO skin samples when compared to WT and *Gnas* E1+/- unaffected skin samples (n=4 samples per condition). (B-C) 12-month dorsal skin sections from (B) WT and (C) *Gnas* E1+/- mice for ⍺SMA (green) and Gli1 (red). (D-E) 12-month dorsal skin sections from (D) WT and (E) *Gnas* E1+/- mice stained for Tgfβ1 (red) and calcein mineral label (green)

### Global deletion of *Sfrp2* within *Gnas E1+/-* mice accelerates SCO formation

Finally, we investigated the functional role of SFRP2 in SCO development based on our initial findings showing increased *SFRP2* expression in fibroblasts isolated from AHO patients, which we also observed in our mouse model. We first carried out cell culture studies using mouse dermal explants (Supplemental Figure 10). Although we observed increased collagen matrix deposition in *Gnas E1+/-* explants compared to *WT*, addition of recombinant SFRP2 to these cultures had no effect on matrix deposition, mineralization, osteogenic differentiation, or *Alpl, Sp7 (Osterix*), and *Ibsp* gene expression.

We next investigated the role of Sfrp2 in SCO formation *in vivo* by crossing *Sfrp2-/-* mice [63] with *Gnas E1+/-* mice. *Sfrp2* deletion was confirmed at both the RNA and protein level in *Gnas E1+/-;Sfrp2-/-* and *Sfrp2-/-* mice by RT-PCR (Figure 8A) and immunofluorescence (Supplemental Figure 11). We examined the rate of SCO formation in both male and female *Gnas E1+/-* and *Gnas E1+/-;Sfrp2-/-* mice by serial radiographic imaging every 4 weeks starting at 4 months (16 weeks) of age. As expected, we did not observe SCO formation in male or female *WT* or *Sfrp2-/-* mice (data not shown); however, we could readily detect SCOs in both *Gnas E1+/-* and *Gnas E1+/-;Sfrp2-/-* mice (Figure 8B-C). As described previously, *Gnas E1+/-* female mice developed fewer SCOs compared to male mice [34] and we did not observe any significant differences in SCO formation between female *Gnas E1+/-* and *Gnas E1+/-;Sfrp2-/-* mice (Supplemental Figure 12). However, male *Gnas E1+/-;Sfrp2-/-* mice developed SCOs significantly earlier than *Gnas E1+/-* mice and also developed a greater number of total SCOs by 20 weeks of age that continued to increase at each subsequent timepoint (Figure 8C). In alignment with their accelerated rate of SCO formation, we observed that *Gnas E1+/-;Sfrp2-/-* mice displayed an expansion in the number of αSMA+ populations throughout the dorsal skin and along the basal epithelial surface when compared to *Gnas E1+/-* mice (Figure 8D,E). The expanded αSMA+ populations in *Gnas E1+/-;Sfrp2-/-* were observed within the basal epithelial surface as well as within epithelial-derived hair follicle populations when compared to *Gnas E1+/-* αSMA+ populations that were limited to the dermis (Figure 8D,E). Collectively, these data demonstrate that loss of Sfrp2 exacerbates the development of SCO formation in *Gnas E1+/-* mice and suggests that *SFRP2* upregulation in AHO may be a compensatory mechanism that limits SCO formation.

**Figure 8:**
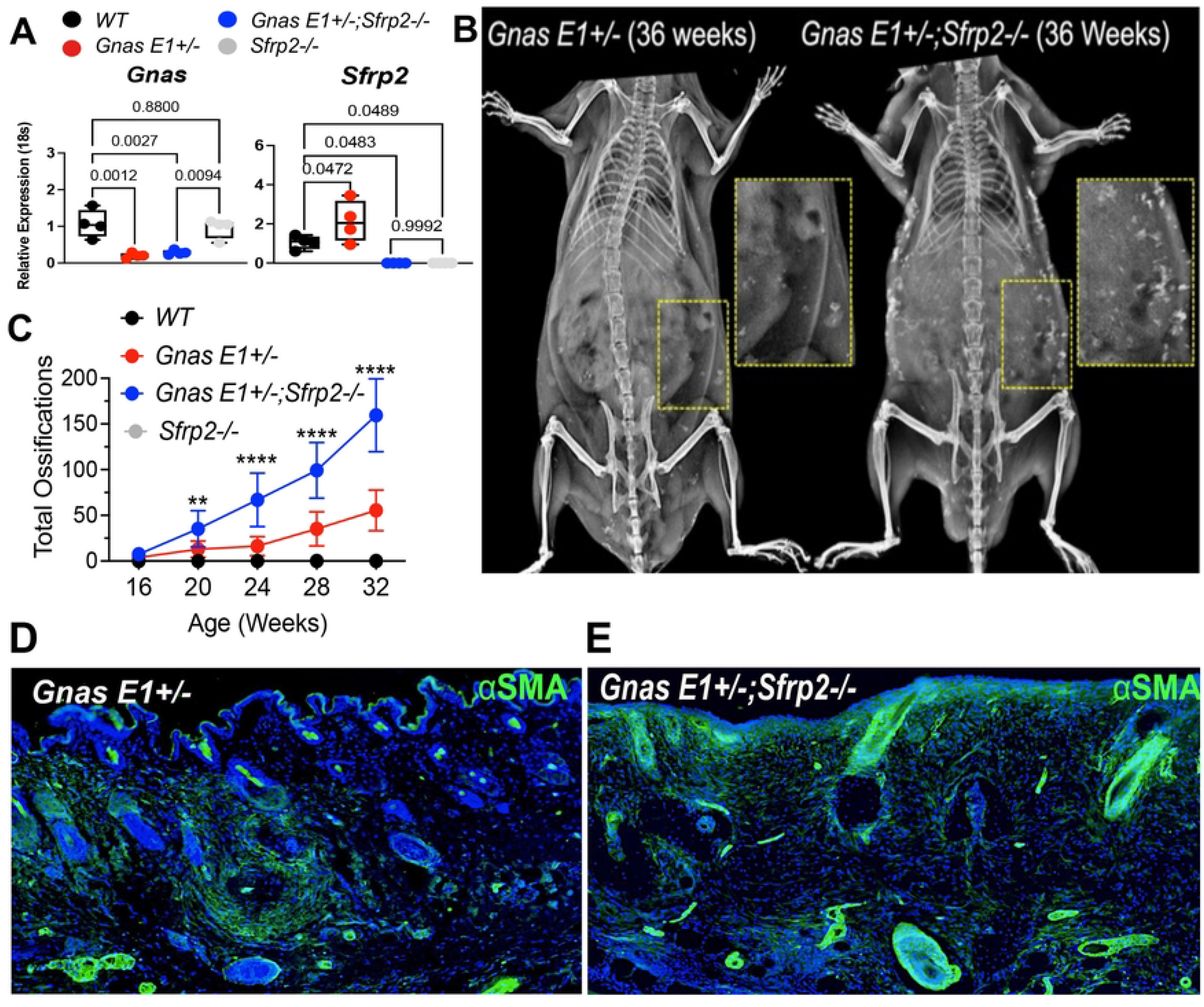
Global deletion of *Sfrp2* within male *Gnas E1+/-* mice accelerates SCO formation. (A) RT-PCR of *Gnas* and *Sfrp2* mRNA expression within 6-month *WT, Gnas E1+/-, Gnas E1+/-;Sfrp2-/-* and *Sfrp2-/-* skin samples. (B) Representative x-ray of 36-week old male *Gnas E1+/-* and *Gnas E1+/-;Sfrp2-/-* mice demonstrating SCO formation. (C) Quantification of SCOs within *Gnas E1+/-* and *Gnas E1+/-;Sfrp2-/-* mice following serial x-ray imaging. (** indicates *p<*0.01, and **** indicates *p*<0.0001). (D-E) Dorsal skin of SCO skin regions of 6-month *Gnas E1+/-* (D) and *Gnas E1+/-;Sfrp2-/-*. (E) mice stained with αSMA (green).

## Discussion

Our data are the first to provide direct evidence that SCO formation in *Gnas E1+/-* mice is initiated by the inappropriate expansion and differentiation of HF-resident dermal sheath cells into osteoblasts. We also demonstrate that following their initial development, SCOs progressively expand through the recruitment and differentiation of a heterogenous population of basal epithelial cells, reactive myofibroblasts, and perivascular cells within the surrounding dermis. These findings correlate with clinical observations in patients with AHO in whom SCOs develop within the dermis and subcutaneous tissue and do not penetrate underlying fascial planes.

We performed lineage tracing using an *Osterix-mCherry* transgenic reporter system [49] in order to characterize the presence of dermal-resident osteoprogenitors. Osterix has been recognized as an indispensable transcription factor necessary for the commitment of mesenchymal progenitors to the osteogenic lineage and to osteoblast differentiation. Our investigations led to the identification of Osterix*+* cell types within the HF of both *WT* and *Gnas E1+/-* mice. These data align with prior studies demonstrating postnatal extraskeletal Osterix expression in tissues such as olfactory glomerular cells [64], intestinal crypt stem cells [64,65] and the renal proximal convoluted tubules [49]. Given that the HF contains multiple epithelial and mesenchymal-derived progenitors, we performed immunofluorescence co-localization studies and identified that these Osterix+ cells specifically corresponded to mesenchymal-derived dermal sheath cells in the HF based on the co-expression studies with αSMA.

The dermal sheath has recently been recognized as a substantial source of mesenchymal progenitors within the HF that exhibit high plasticity and self-renewal capacity throughout adulthood (for review, see [66]). *In vivo* fate-mapping studies of dermal sheath cells using a tamoxifen-inducible *⍺SMACreER^T2^;YFP* mouse model have demonstrated that there is a subset of HF stem cells within the dermal sheath that can regenerate out to 24 months post tamoxifen injection and can differentiate and repopulate into multiple cell types within the regenerating HF [50,67]. In addition, studies using human and rat-derived dermal sheath cells have been shown to differentiate into adipocytes, osteoblasts and chondrocytes *in vitro* [36,68]. These data, in conjunction with our genetic fate-mapping studies demonstrating that *Gnas E1+/-* dermal sheath cells differentiate into osteoblasts and osteocytes, further emphasize the plasticity of these HF-resident mesenchymal progenitors. Future transplantation studies assessing the osteogenic capacity of purified dermal sheath cells using methods such as a subcutaneous implantation or calvarial defect models are warranted.

Our genetic fate-mapping studies in 10-month-old *Gnas E1+/-;Osx-mCherry* mice already containing SCOs identified Osterix+ populations dispersed throughout the surrounding epidermis and dermis. We hypothesize that this heterogenous expression pattern within the dermis is attributed to the recruitment of activated dermal myofibroblasts and pericytes in response to lesion development. In addition, our observations of Osterix expression within basal epithelial cells and epithelial-derived HF progenitors implicate that these populations are undergoing epithelial-mesenchymal transition. These observations directly correlate with prior fate-mapping studies that have shown that in postnatal *Gnas* homozygous deletion models, initial heterotopic bone formation is driven by the osteogenic differentiation of local mesenchymal progenitors; however, the progressive expansion of these lesions over time is driven by the recruitment of surrounding cell types to the lesion site for subsequent osteogenic differentiation [69,70].

In conjunction with assessing the cellular populations that contribute to SCO formation, we also identified significant variations in Sonic Hedgehog, TGF-β, BMP, and Wnt signaling activity within SCO-containing skin regions of *Gnas E1+/-* mice by RT-PCR array and immunofluorescence. These data are of particular interest given that each of these pathways have been shown to contribute to the pathogenesis of trauma-induced HOs [1,71–75] but to date have not been fully implicated in the context of ossification formation in AHO or POH. Consequently, further studies are warranted to further understand the role of the Gα_s_-PKA-cAMP signaling cascade in mediating epithelial-mesenchymal interactions within the hair follicle microenvironment. Additionally, despite these encouraging data, it is also important to acknowledge that these findings are representative of the global transcriptional changes within the skin at 12 months of age with the presence of established SCOs contained within the dermis. To this point, it is likely that there are additional and alternative contributing pathways throughout the various stages of SCO development that are restricted to specific timeframes, tissue microenvironments and/or individual cell populations that may not be most representative within this current data set. Consequently, these data pose an interesting set of additional questions into the spatiotemporal activation of varying cell types involved in the pathogenesis of heterotopic bone formation that warrants further exploration though the use of techniques such as single cell RNA sequencing and spatial transcriptomics.

Finally, we had carried out microarray analysis of RNA from dermal fibroblast cultures derived from skin biopsies from patients with AHO and POH presenting with varying degrees of SCOs. By comparing gene expression profiles from these patients, we had identified a direct correlation between *SFRP2* mRNA expression and SCO severity. Although SFRP2 was initially characterized as a secreted protein that competitively binds and inhibits ligands essential for canonical Wnt signaling [39–41,43,76], more recent studies have identified this gene to function as a negative regulator of both epithelial proliferation and epithelial to mesenchymal transition [77–79]. We also analyzed Sfrp2 expression in the skin of *Gnas E1+/-* mice and similarly found strong expression within the HF and its surrounding dermis and basal epithelium. Our findings could be consistent either with SFRP2 playing a causal role in the initiation of SCO development or with SFRP2 upregulation being a response to mitigate further SCO formation. In order to distinguish these two possibilities, we investigated the development of SCOs in *Gnas E1+/-;Sfrp2-/-* mice. We found that loss of *Sfrp2* exacerbated the development of SCOs resulting from heterozygous loss of *Gnas* in terms of both age of onset and total number of SCOs. These results imply that the upregulation of *SFRP2* seen in both humans and mice likely represents a compensatory mechanism that limits further SCO development and/or progression. Our findings raise the possibility that administration of SFRP2 or a functional agonist may be a potential therapeutic strategy for the prevention and treatment of heterotopic ossifications.

## Materials and Methods

### Generation and maintenance of mice

All animal studies and protocols were carried out in accordance with the standards of the UConn Health Animal Care and Use Committee. Mice were fed a standard diet of mouse chow and water *ad libitum*. Mouse strains and their background are outlined in Table 1. The generation of *Gnas E1+/-* mice carrying a targeted disruption of exon 1 of *Gnas* [27,34] as well as the generation and characterization of *Osx-mCherry*[49] mice and *⍺SMACre^ERT2^;Ai9^fl/fl^* [53] mice have been previously described. *Sfrp2^tm1.1Brle^* mice (from here on termed *Sfrp2-/-* mice) were purchased from Jackson Labs. *Gnas E1+/-, Sfrp2-/-* and *⍺SMACre^ERT2^;Ai9^fl/fl^* mice were maintained on a pure 129SvEv background. *Osx-mCherry* mice were maintained on a CD1 background; therefore for *Gnas E1+/-;Osx-mCherry* mice used for fate-mapping studies, the mice were bred as F1 129xCD1 crosses and the *Osx-mCherry* littermates were used as controls. Mice were genotyped by PCR analysis using the primer sequences outlined in Table 2. For experiments utilizing *⍺SMACre^ERT2^;Ai9^fl/fl^* mice, Cre activation was performed by intraperitoneal injection of tamoxifen in corn oil (75 µg/g body weight). Mice were administered 2 doses of tamoxifen spaced 48 hours apart.

**Table 1:**
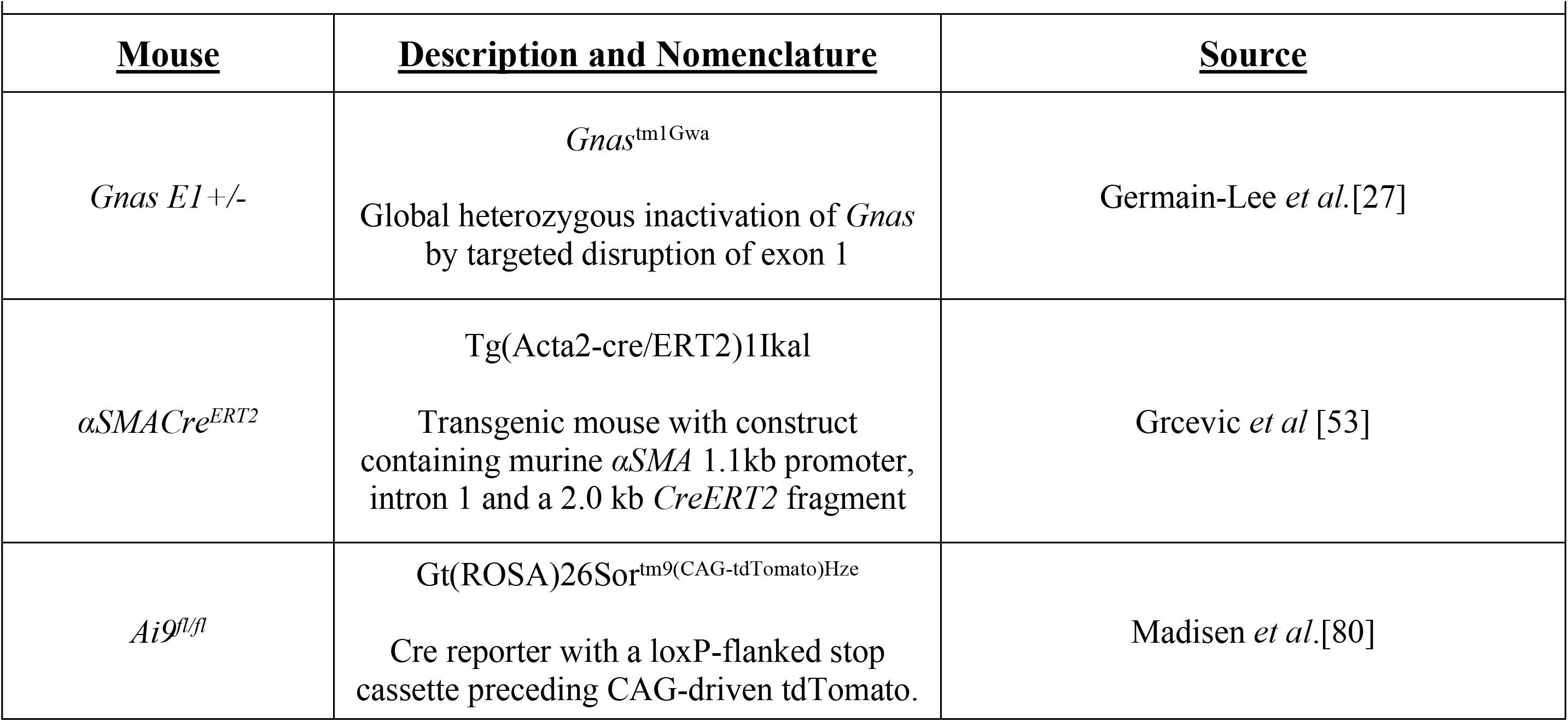
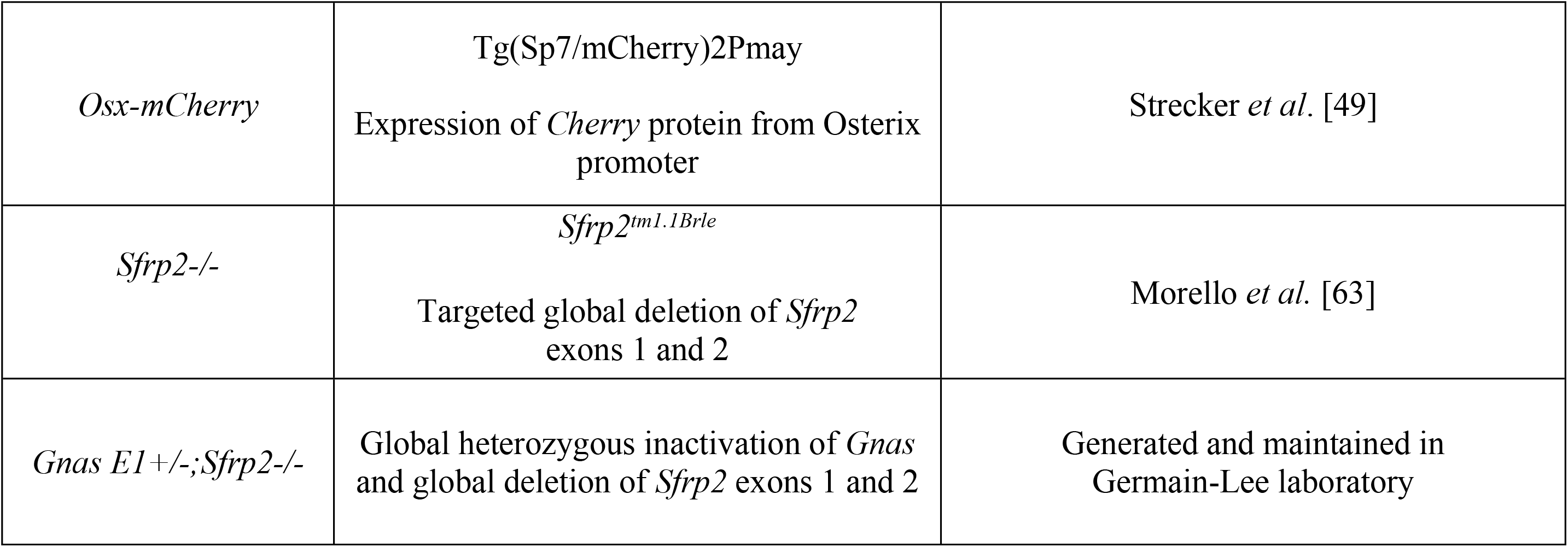
Mouse Models Utilized.

**Table 2:**
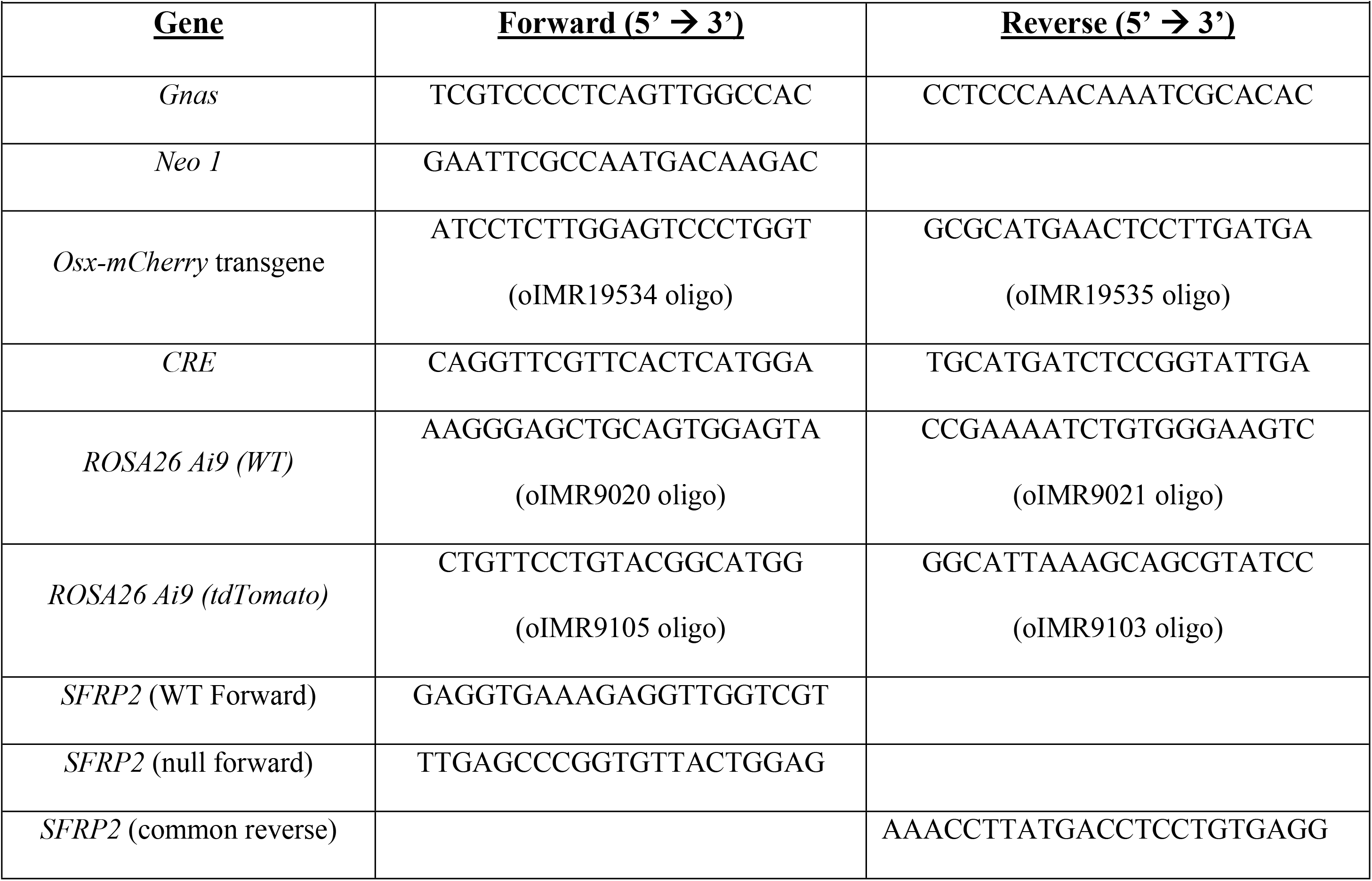
PCR Oligonucleotide Primer Sequences.

### Human GNAS mutation analyses of participants

All human studies were approved by the Johns Hopkins Medicine Institutional Review Board (E.L.G-L.’s institution at that time). Informed consent was obtained from all participants (or parent of participant) prior to enrollment. Assent was obtained when appropriate based on age and emotional/cognitive maturity. In brief, peripheral blood from all participants was collected within the Johns Hopkins Institute of Clinical and Translational Research. DNA isolation and *GNAS* mutation analyses of the 13 coding exons and all intron/exon boundaries, including determination of the parental origin of the mutated allele, were performed for all participants in our investigations (E.L.G-L., Johns Hopkins laboratory) as previously described [22,24,25,81]. Therefore, all participants were mutation-confirmed. We also documented that the POH participant in our study had inheritance of the mutation from the paternal allele, which is the pattern of inheritance typical of POH [81]. The participant with POH had very severe SCOs as well as deep, penetrating ossifications assessed by surgical pathology of excised ossifications as well as CT scan and magnetic resonance imaging (MRI) performed for clinical reasons.

### Human dermal biopsy culture generation from mutation-confirmed AHO and POH participants

Informed consent and assent specifically for the skin biopsies was additionally obtained from all participants (or parent of participant) prior to enrollment. For consistency, skin biopsies were performed by the same investigator (E.L. G-L.) on all participants with mutation-confirmed AHO (both PHP1A and PPHP) and POH, as well as on unaffected family members (in whom no *GNAS* mutations were identified). Skin biopsies were performed according to standard clinical procedures using a 2 mm biopsy punch (Accu-Punch Biopsy Punch, Accuderm Inc, Ft Lauderdale, FLA) after numbing the region with 1% lidocaine. Biopsies of SCO regions were performed on an extremity within a location of significant subcutaneous tissue. If no SCOs were present, biopsies were collected on the ventral surface of the participant’s forearm. Primary cultures were generated and maintained in Minimum Essential Medium (Eagle’s) with Earle’s Salts supplemented with 85 units of Penicillin and Streptomycin/ml, 1.7 moles L-glutamine and 13% Fetal Bovine Serum (FBS). RNA was extracted using a cesium chloride gradient according to routine well-established procedures as previously described [82]. There were no complications post-procedure.

### Microarray analyses from RNA generated from human dermal cultures

Microarray analyses were performed using RNA obtained from dermal fibroblast cultures from 5 mutation-confirmed participants, consisting of 4 with AHO and 1 with POH. Participants with AHO with varying degrees of SCOs were chosen. The severity of SCOs was based on the number of individual lesions noted on palpation by one consistent examiner (E.L.G-L.) and categorized as either none, minimal, moderate, or severe with minimal < 3, moderate = 3 - 25, and severe > 25. The number of SCOs > 1.0 cm increased with the degree of severity.

All microarray analyses were performed by the Johns Hopkins Medical Institutions Microarray Core Facility using an Affymetric GeneChips U133 Plus 2.0 (human) chip (Santa Clara, CA). Differential gene expression was determined on the basis of exhibiting either > +2.0 fold or < -2.0 fold change as well as a having a probability of > 0.50. Pairwise comparisons of these samples (ie. two participants analyzed and compared to one another) were performed as shown in Figure 1A. In all three comparisons (P1 x P2, P3 x P2, and P4 x P5), a participant with the greater number (as well as size) of SCOs was compared to a participant with fewer or no ossifications. The distribution of the final subset of differentially expressed genes is shown in Figure 1, panels B and C.

### Human SFRP2 northern blot studies

RNA isolation and Northern blot analysis from human cultured fibroblasts were performed as previously described[82] using 10 micrograms of RNA per lane and analyzed by phosphorimager quantitation (Bio-Rad, Hercules, CA) of the resulting autoradiograph of the Northern blot (Bio-Rad, Hercules, CA) and expressed as a ratio of *SFRP2* to *S26* [82]. For these studies, there were an additional 2 participants with mutation-confirmed AHO, and therefore a total of 9 participants: 6 with AHO (2 with PHP1A and 4 with PPHP), 1 with POH, and 2 unaffected family members for whom sequencing revealed no mutation in *GNAS* (Figure 1D).

### Mouse histology

Following euthanasia by CO2 asphyxiation, dorsal skin hair was removed using an electric trimmer and a depilatory cream (Nair, Church & Dwight, New York, NY). Dorsal skin samples, including the underlying adipose and muscle were harvested and fixed in 10% neutral buffered formalin (NBF) overnight at 4°C, followed by 30% sucrose in PBS for 24 hours at 4°C and subsequently embedded into Optimal Cutting Temperature (OCT). Tissue blocks were stored at -20°C until use, and cryosections (10-15μm) were collected using a cryostat tape-transfer system (Section-lab, Hiroshima, Japan) as previously described [83]. Samples were imaged using a Ziess Axioscan Z1 high-speed automated image acquisition system (Cat#440640-9903-000) and a high resolution camera (AxioCam HRm).

### Bone mineral label visualization

*WT* and *Gnas E1+/-* mice were administered intraperitoneal injections of calcein [10mg/kg (Sigma C-0875)] or alizarin complexone (30mg/kg (Sigma A-3882)] 2-7 days prior to sacrifice. Dorsal skin sections were stained with calcein blue (Sigma M1255) for 10 minutes to visualize total mineral content, as previously described [30,83]. For multiplex staining, skin sections were decalcified using a sodium acetate and sodium tartrate dibasic dihydrate solution in water (pH 4.2) to remove bone mineral labels from the tissue section as described previously [83].

### Immunofluorescence

Antibodies utilized for immunofluorescence staining are listed in Table 3. Tissues were permeabilized in 0.1% Triton X for 10 minutes, blocked using a Mouse-on-Mouse Immunodetection Kit (Vector Laboratories BMK-2202) diluted in a 2% bovine serum albumin in PBS solution for 45 minutes and stained with primary antibody overnight at 4°C. The following day, sections were washed in 0.1% Triton X in PBS for 10 minutes and stained with secondary antibodies (1:300 dilution) in 2% BSA in PBS at room temperature for 1 hour. Tissue sections were mounted in a 50% glycerol / PBS solution containing 4’,6-diamidino-2-phenylindole (DAPI) (1:1000 dilution) and imaged.

**Table 3:**
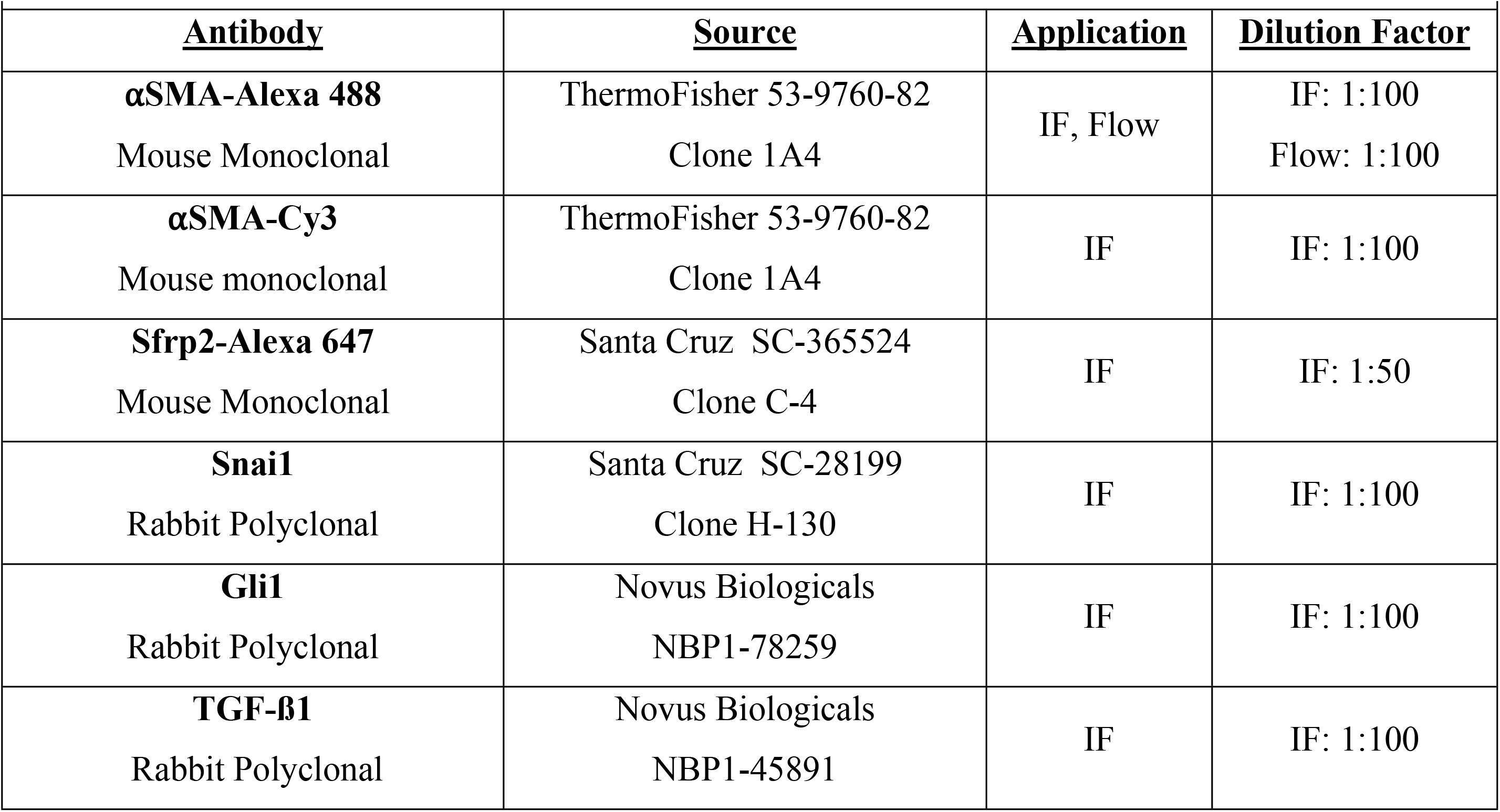
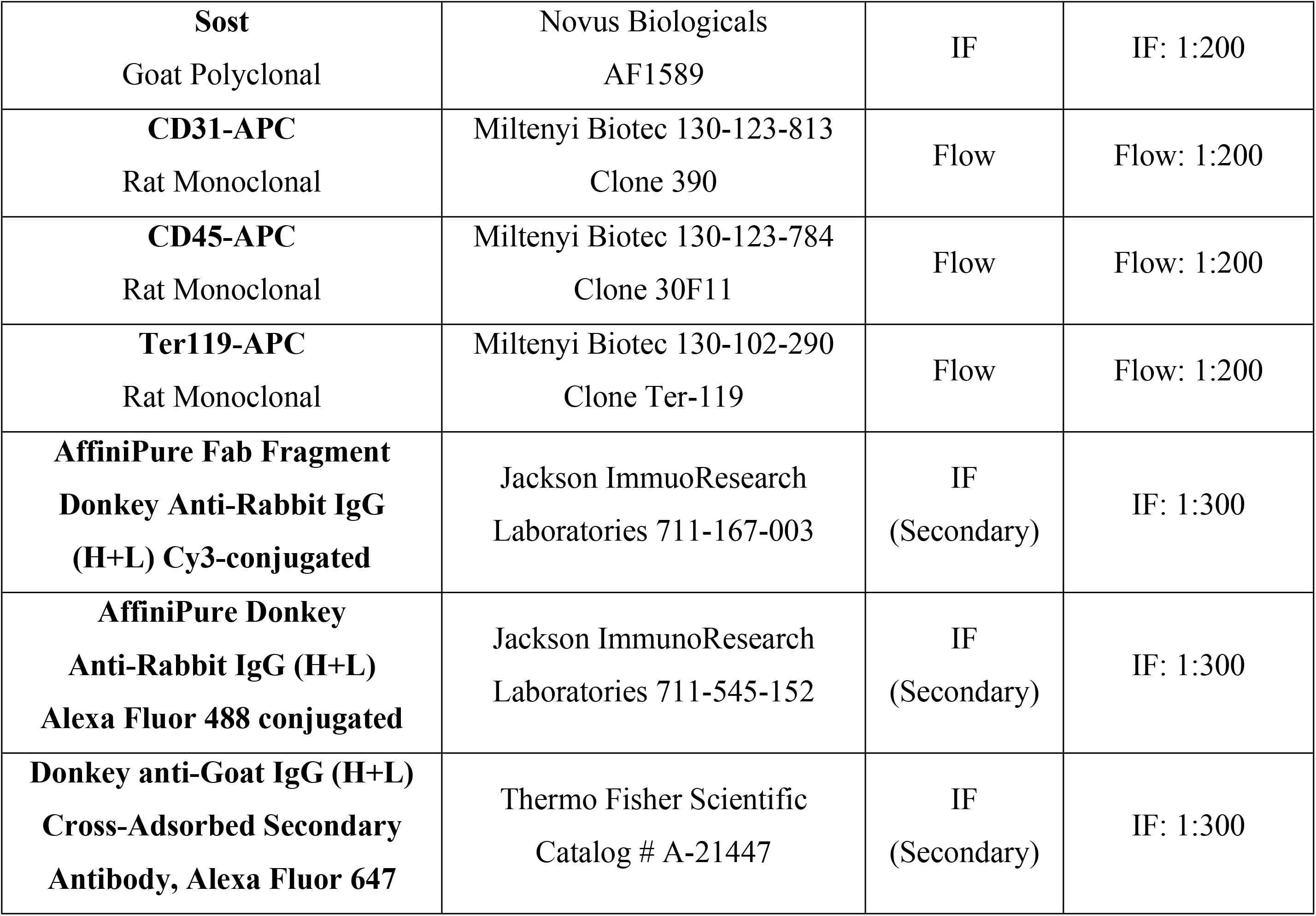
Antibodies Utilized for Immunofluorescence and Flow Cytometry.

### ALP enzyme histochemistry

Alkaline phosphatase (ALP) enzyme histochemistry was performed on tissue sections following the methods previously described [30,83,84]. Briefly, slides were incubated in an alkaline buffer (1M Tris, 1M MgCl_2_, 2 M NaCl in deionized water pH 9.5) for 10 minutes, followed by exposure to an alkaline Fast Red substrate buffer for 30 minutes. Sections were mounted in 50% glycerol/PBS solution with DAPI and coverslipped for image acquisition. For experiments performed with *Osx-mCherry* and *⍺SMA-CRE^ERT2^;Ai9^fl/fl^* reporter mice, ALP staining was performed using a Vector Blue Alkaline Phosphatase Substrate Kit (Vector Laboratories SK-5300) for 30 minutes to avoid overlap of fluorescent signals.

### Chromogenic staining

General tissue architecture within the dermis was visualized by staining tissues with 0.025% toluidine blue in deionized water for 5 minutes as previously described [83]. Von Kossa staining was performed by incubating slides with 4% silver nitrate solution and exposing slides to 2400 kJ of ultraviolet light using a UV Stratalinker. Safranin O and fast green staining were performed by staining sections in Weigert’s iron hematoxylin for 5 minutes to visualize nuclei, rinsed in tap water for 10 minutes, stained in 0.2% fast green solution for 2 minutes, rinsed in 1% acetic acid solution for 2 minutes, and stained in 0.1% safranin O solution for 1 minute. Masson trichrome staining was performed using a commercially available kit (Sigma HT15-1KT) based on the manufacturer’s instructions.

### Dorsal Skin Flow Cytometry

Flow cytometry analyses were performed using single cell suspensions isolated by enzymatic digestion from 10 month old *WT* and *Gnas E1+/-* dorsal skin samples using slightly modified methods as described by *Walmsley* et al, (2016) [85]. Briefly, the entire dorsal skin was harvested, avoiding collection of underlying adipose or skeletal muscle, and placed in sterile PBS on ice. Harvested samples were minced into 1-2 mm fragments and placed into DMEM supplemented with 2mg/mL collagenase IV, and 0.5 mg/mL collagenase I. Collection beakers were then placed onto a magnetic stirrer at medium speed inside a cell culture incubator at 37°C to promote tissue digestion. After 90 minutes of digestion, an equal amount of DMEM containing 10% FBS was added to the digested tissue sample, passed through a 100 µm filter, and centrifuged at 300 g for 10 minutes at 4°C. The collected cell pellet was resuspended in Zombie Fixable Live/Dead Staining Solution in PBS and incubated for 30 minutes at 4°C protected from light. Samples were subsequently washed and centrifuged in Staining Media [(Hanks’ Balanced Salt Solution (HBSS) supplemented with 10% FBS and 10µg/mL DNase I)]. The remaining pellet was stained for 20 minutes at 4°C in Staining Media with primary conjugated antibodies for cell surface markers (antibodies, clones, and dilution factors are summarized in Table 4). Stained cells were washed, centrifuged, fixed in 4% PFA for 15 minutes at 4 °C and incubated in 1X InVitrogen eBioscience Cell Permeabilization buffer (InVitrogen 00-8333-56) for 15 minutes at 4 °C. After centrifugation, the resuspended pellet was stained with an Alexa-488 conjugated ⍺SMA antibody clone 1A4 (1:100 dilution) for 30 minutes at 4 °C. Washed and filtered cells were analyzed with a BD-LSRII flow cytometer with gates established according to unstained and single-channel controls. All compensations were identified using FACS Diva Software, and downstream data analysis was performed using FlowJo Software v10.

**Table 4.**
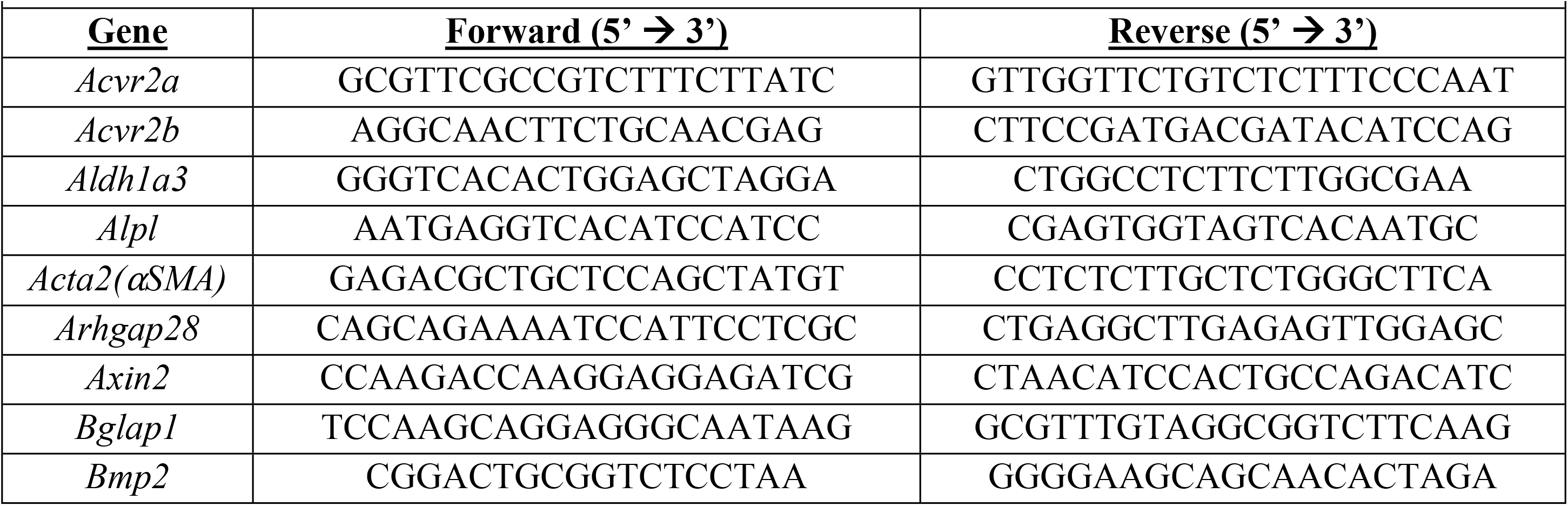
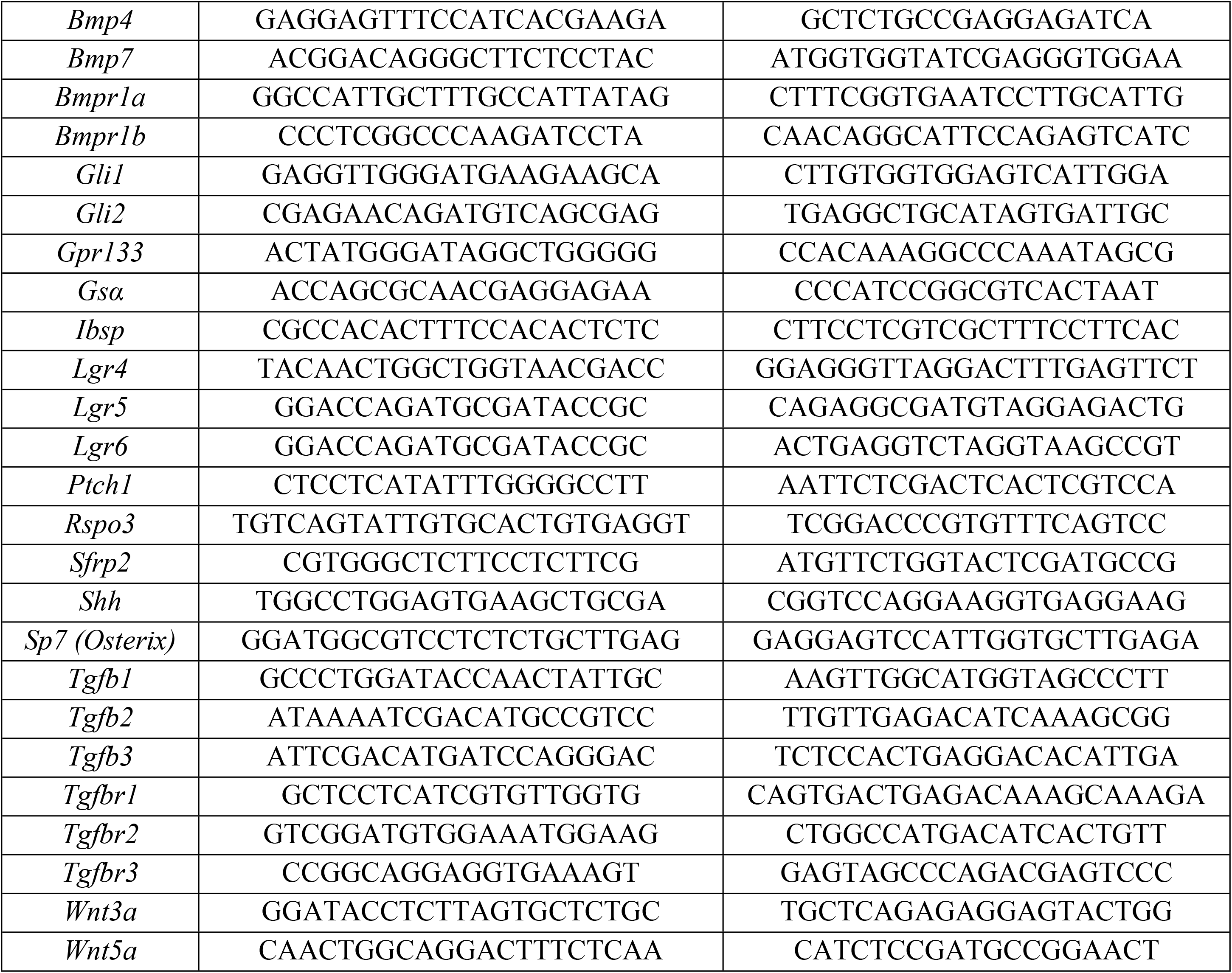
Real-time Quantitative RT-PCR Oligonucleotide Primer Sequences. (Note: All Forward and Reverse Primer Sequences are Located on Different Exons)

### Primary dermal explant cultures

Primary dermal explant cultures were established from dorsal skin fragments isolated from 9-month *Gnas E1+/-;⍺SMA-CRE^ERT2^;Ai9^fl/fl^* and *⍺SMA-CRE^ERT2^;Ai9^fl/fl^* mice following methods as previously described by *Seluanov et al* 2010 [57]. Confluent primary cell cultures were dissociated using Accutase and subsequently prepared for either fluorescence activated cell sorting (FACS) or passaged onto 24-well culture dishes at a cellular density of 5.0×10^4^ cells per well for osteogenic differentiation or collagen deposition assays. For experiments using recombinant Sfrp2 treatment, cultures were exposed for the duration of treatment with either 100ng/mL recombinant mouse Sfrp2 (R&D Systems Cat 1169-FR-025) or vehicle control (PBS with 0.1% BSA).

### Dermal explant FACS sorting

FACS analysis and cell sorting of Ai9+ cells from primary dermal explants were performed using a BD FACSAria II (BD Biosciences, San Jose, CA, USA). Primary cell culture pellets were resuspended into FACS Staining Media (HBSS supplemented with 10% FBS) containing Sytox Blue Dead Cell Stain (InVitrogen S34857). Sorting gates were established using cultures generated from non-transgenic cultures and unstained transgenic cultures. All downstream data analysis was performed using FlowJo in order to determine the percentage of Ai9+ sorted populations.

### In vitro osteogenic differentiation assays

Osteogenic differentiation capacity was assessed *in vitro* by exposing cultures to DMEM/F12 with 10% FBS, 1% penicillin/streptomycin supplemented with 50μg/mL ascorbic acid and 10mM Beta-glycerophosphate for 28 days. Live imaging studies assessing bone mineralization was performed by supplementing culture media with 30μM calcein (Sigma C-0875) overnight. The following day, areas of mineralization and their localization to Ai9+ populations were assessed by fluorescence microscopy. Cultures were subsequently fixed in 4% PFA and stained for ALP using the Vector Blue Alkaline Phosphatase Substrate Kit (Vector Laboratories SK-5300) and Von Kossa using a 4% silver nitrate solution and exposing cultures to 2400 kJ of ultraviolet light using a UV Stratalinker.

### In vitro collagen deposition assays

Total collagen deposition from dermal explants cultured in DMEM/F12 containing 10% FBS for 14 days was assessed using a Sirius red/fast green assay as previously described [86,87]. Briefly, cells were fixed using a Kahle fixative solution, stained for 30 minutes in a 0.1% Fast Green and 0.2% Sirius Red solution in saturated picric acid in distilled water. Stained cultures were visualized under brightfield microscopy to assess collagen deposition and destained using a 0.1N sodium hydroxide solution in absolute methanol for absorbance measurements at 540nm and 605nm. Calculations of total collagen per well using these absorbance measurements were performed based on the previously described methods [87].

### RNA purification

Total RNA was extracted from dermal explant cultures, FACS-sorted cultures and 1cm^2^ dorsal skin samples using an RNEasy Micro Kit (Qiagen) for FACS-sorted cultures and a Direct-zol RNA Miniprep Kit (Zymo Research) for dermal explant and dorsal skin samples following the manufacturer’s instructions. Prior to RNA isolation, harvested dorsal skin tissue was placed into RNA Later solution (InVitrogen) and stored at -80°C. Prior to RNA isolation, the tissue was thawed into RNA Later solution, minced into 1-2mm fragments and subsequently placed into 1mL of Trizol (Invitrogen) on ice and homogenized. RNA samples were treated with DNAse I (New England Biosciences) and were concentrated through a Monarch RNA Cleanup Kit (NEB) to ensure no carryover of contaminants.

### Quantitative RT-PCR

1 μg of RNA was utilized for reverse transcription using a high capacity cDNA reverse transcription kit (Applied Biosystems). Quantitative RT-PCR was performed using a Bio-Rad CFX96 ThermoCycler (Bio-Rad Laboratories, Hercules, CA) within a 20 μL reaction, consisting of iTaq Universal SYBR Green supermix (Bio-Rad Laboratories, Hercules, CA), 10 μM of forward and reverse primers and 25ng of cDNA. The specific primer sequences utilized are listed in Table 4.

### Statistical analysis

All statistical analyses were performed using Graphpad Prism Version 9 (GraphPad Software, Inc., La Jolla, CA, USA) with *P*-values < 0.05 considered statistically significant. For all analyses that compared data obtained from *WT* and *Gnas E1+/-* mice at one discrete time point (i.e. Flow and FACS analyses and cell culture RT-PCR studies), data were analyzed using an unpaired two-tailed t-test. For all analyses observed at one discrete timepoint comparing data from three or more groups from *WT, Gnas E1+/-, Sfrp2-/-* or *Gnas E1+/-;Sfrp2-/-* mice, the data were analyzed by a one-way ANOVA with a post-hoc Tukey test for multiple comparisons. Each *n* value refers to the number of mice. All data points were included in the data analysis. No samples were excluded. *P*-values for each statistical comparison are indicated within the figure panels.

All mouse analyses and human data are included within the main and supplemental figures within this manuscript.

## Authors CRediT Contribution Roles

Patrick McMullan – Conceptualization, Data Curation, Formal Analysis, Investigation, Methodology, Validation, Visualization, Writing – Original Draft Preparation, Writing – Review, Writing – Editing

Peter Maye – Methodology, Project Administration, Resources, Supervision, Validation, Writing

– Review, Writing – Editing

Sierra Root - Methodology, Data Curation, Formal Analysis, Investigation, Validation, Visualization, Writing – Review, Writing – Editing

Qingfen Yang – Data Curation, Investigation, Methodology, Validation, Writing – Review Sarah Edie – Data Curation, Investigation, Validation, Writing – Review, Writing – Editing David W. Rowe – Methodology, Resources, Supervision, Validation, Writing – Review

Ivo Kalajzic - Methodology, Resources, Supervision, Validation, Writing – Review, Writing – Editing

Emily L. Germain-Lee – Conceptualization, Data Curation, Formal Analysis, Funding Acquisition, Investigation, Methodology, Project Administration, Resources, Supervision, Validation, Writing – Original Draft Preparation, Writing – Review, Writing – Editing

## Acknowledgments

We sincerely thank the patients and their families who made this work possible. We also thank the Johns Hopkins University School of Medicine Institute of Clinical and Translational Research and the Johns Hopkins Microarray Core Facility, as well as Evan Jellison and the UConn Health Flow Cytometry Core Facility for their assistance in a portion of these studies. The authors declare that there are no conflicts of interest.

## Authors’ roles

P.Mc., P.Ma., S.R., I.K., D.R., and E.L.G-L. contributed to study design. P.Mc., S.R., S. E., Q.Y., and E.L.G-L. collected and analyzed data. P.Mc. and E.L.G-L. wrote the initial manuscript. P.Mc., P.Ma., S.R., S.E., I.K., and E.L.G-L. critically revised the manuscript.

---

**Supplemental Figure 1:** Quantitative RT-PCR analysis of upregulated genes identified through AHO/POH human fibroblast microarray comparisons (*Arhgap28, Aldh1a3, Gpr133* and *Ror2*) within 12-month *WT* and *Gnas E1+/-* skin samples. Statistical analyses were completed using ANOVA with post-hoc Tukey test for multiple comparisons. *P*-values for each statistical comparison are indicated within the panels.

**Supplemental Figure 2:** Dorsal skin section of 15-month *Gnas E1+/-* mouse isolated from SCO region and stained with αSMA (red) and Sfrp2 (yellow).

**Supplemental Figure 3:** Representative images of dorsal skin sections isolated from 10-month *Gnas E1+/-* mice following: (A) Masson Trichome; (B) Safranin O/Fast Green; and (C) Tartrate resistant acid phosphatase (TRAP)/Aniline Blue staining.

**Supplemental Figure 4:** (A) Schematics of transgene and breeding strategies. (B) Representative images of dorsal skin sections isolated from 3-week old (P21) males: (top panel) *Osx-mCherry* and (bottom panel) *Gnas E1+/-;Osx-mCherry* littermates demonstrating similar expression patterns of *Osterix+* populations for the hair follicle dermal sheath.

**Supplemental Figure 5:** Flow cytometry gating strategy implemented to quantify the percentage of ⍺SMA+ cells within the Lineage-negative population from dorsal skin single cell suspensions as depicted in Figure 3.

**Supplemental Figure 6:** Representative images of dorsal skin section from 10-month *⍺SMACre^ERT2^;Ai9^fl/fl^* and *Gnas E1+/-;⍺SMACre^ERT2^;Ai9^fl/fl^* mice in both unaffected and SCO skin regions without tamoxifen administration.

**Supplemental Figure 7:** Representative images of dorsal skin sections from 2-month old *⍺SMACre^ERT2^;Ai9^fl/fl^* and *Gnas E1+/-;⍺SMACre^ERT2^;Ai9^fl/fl^* mice at 2 days post tamoxifen injection. Note the visualization of Ai9+ populations within the hair follicle dermal sheath, hair follicle arrector pili muscles, and underlying blood vasculature.

**Supplemental Figure 8:** Unstained and single color staining controls utilized to establish FACS sorting gates for *⍺SMACre^ERT2^;Ai9^fl/fl^* and *Gnas E1+/-;⍺SMACre^ERT2^;Ai9^fl/fl^* dermal explant cultures. Additionally, aliquots of sorted tdTomato- and Ai9+ populations were reanalyzed to assess purity.

**Supplemental Figure 4:** (A) Schematics of transgene and breeding strategies. (B) Representative images of dorsal skin sections isolated from 3-week old (P21) males: (top panel) *Osx-mCherry* and (bottom panel) *Gnas E1+/-;Osx-mCherry* littermates demonstrating similar expression patterns of *Osterix+* populations for the hair follicle dermal sheath.

**Supplemental Figure 5:** Flow cytometry gating strategy implemented to quantify the percentage of ⍺SMA+ cells within the Lineage-negative population from dorsal skin single cell suspensions as depicted in Figure 3.

**Supplemental Figure 6:** Representative images of dorsal skin section from 10-month *⍺SMACre^ERT2^;Ai9^fl/fl^* and *Gnas E1+/-;⍺SMACre^ERT2^;Ai9^fl/fl^* mice in both unaffected and SCO skin regions without tamoxifen administration.

**Supplemental Figure 7:** Representative images of dorsal skin sections from 2-month old *⍺SMACre^ERT2^;Ai9^fl/fl^* and *Gnas E1+/-;⍺SMACre^ERT2^;Ai9^fl/fl^* mice at 2 days post tamoxifen injection. Note the visualization of Ai9+ populations within the hair follicle dermal sheath, hair follicle arrector pili muscles, and underlying blood vasculature.

**Supplemental Figure 8:** Unstained and single color staining controls utilized to establish FACS sorting gates for *⍺SMACre^ERT2^;Ai9^fl/fl^* and *Gnas E1+/-;⍺SMACre^ERT2^;Ai9^fl/fl^* dermal explant cultures. Additionally, aliquots of sorted tdTomato- and Ai9+ populations were reanalyzed to assess purity.

**Supplemental Figure 9:** Quantitative RT-PCR analysis of Sonic Hedgehog, TGF-beta, BMP and Wnt signaling pathway transcripts analyzed among 12 month *WT* and *Gnas E1+/-* skin samples as depicted in the heat map in Figure 7. Statistical analyses were completed using ANOVA with post-hoc Tukey test for multiple comparisons. *P*-values for each statistical comparison are indicated within the panels.

**Supplemental Figure 10:** (A) Representative whole well images of Sirius Red staining and quantification of total collagen per well observed in dermal explant cultures following 14 days of exposure to 100ng/mL recombinant mouse Sfrp2 or vehicle control. (B) Quantitative RT-PCR of *Gnas E1+/-* dermal explant cultures following 28 days of osteogenic differentiation media supplemented with 100 ng/mL recombinant mouse Sfrp2 or vehicle control. (C) Statistical analyses for Sirius red staining were completed using ANOVA with post-hoc Tukey test for multiple comparisons; statistical analyses of osteogenic differentiation RT-PCR studies were performed using an unpaired two-sided T test. *P*-values for each statistical comparison are indicated within the panels.

**Supplemental Figure 11**: Representative immunofluorescence staining for αSMA (red) and Sfrp2 (yellow) within dorsal skin sections of 6 month *WT, Gnas E1+/-, Sfrp2-/-* and *Gnas E1+/-;Sfrp2-/-* mice in unaffected skin regions.

**Supplemental Figure 12:** (A) Quantification of total SCOs identified within female *Gnas E1+/-* and *Gnas E1+/-;Sfrp2-/-* mice (B) Quantification of total SCOs identified between male and female *Gnas E1+/-* mice up to 32 weeks of age. (C) Quantification of total SCOs identified between male *Gnas E1+/-* and *Gnas E1+/-;Sfrp2-/-* mice with either maternally-inherited or paternally-inherited *Gnas* mutations.

